# Cooling of medial septum reveals theta phase lag coordination of hippocampal cell assemblies

**DOI:** 10.1101/2019.12.19.883421

**Authors:** Peter C. Petersen, György Buzsáki

**Affiliations:** Neuroscience Institute, Langone Medical Center, New York University, New York, NY 10016, USA; Department of Neurology, Langone Medical Center, New York University, New York, NY 10016, USA; Center for Neural Science, New York University, New York, NY 10003, USA

## Abstract

Hippocampal theta oscillations coordinate neuronal firing to support memory and spatial navigation. The medial septum (MS) is critical in theta generation by two possible mechanisms: either a unitary ‘pacemaker’ timing signal is imposed on the hippocampal system or it may assist in organizing target subcircuits within the phase space of theta oscillations. We used temperature manipulation of the MS to test these models. Cooling of the MS reduced both theta frequency and power, was associated with enhanced incidence of errors in a spatial navigation task but did not affect spatial correlates of neurons. MS cooling decreased theta frequency oscillations of place cells, reduced distance-time compression but preserved distance-phase compression of place field sequences within the theta cycle. Thus, septal computation contributes not only theta pacing but is also critical for sustaining precise theta phase-coordination of cell assemblies in the hippocampus.

## Introduction

Theta frequency oscillations coordinate neuronal activity in the hippocampus-subicular-entorhinal complex and entrain neurons in various neocortical areas (Buzsáki, 2002). Damage or inactivation of the medial septum–diagonal band (MS) abolishes theta oscillations in all these areas (Petsche et al., 1962; Vertes and Kocsis, 1997) and is associated with impairment of memory, spatial navigation and other cognitive functions (Bolding et al., 2020; Brandon et al., 2011; Chang and Gold, 2004; Chrobak et al., 1989; Givens and Olton, 1990; Jeffery et al., 1995; Leutgeb and Mizumori, 1999; Wang et al., 2015; Winson, 1978). However, whether the behavioral impairment is due to silencing or damaging important septal afferents or to the absence of theta phase-multiplexed coordination of activity of neurons (Harris et al., 2003; Kay et al., 2020) in the hippocampal system has remained an unsolved challenge. Such dissociation is not straightforward because manipulations that abolish theta also affect neurons and synapses which may exert their own, theta-independent effects.

Early experiments suggested that the MS acts as a ‘pacemaker’, sending out synchronous outputs, akin to a conductor of an orchestra (Borhegyi, 2004; Petsche et al., 1962; Stewart and Fox, 1990; Sweeney et al., 1992; Zutshi et al., 2018), followed by models in which cholinergic and GABAergic neurons of MS fire at distinct unique phases of the theta cycle (Borhegyi, 2004; Petsche et al., 1962; Stewart and Fox, 1990; Sweeney et al., 1992). However, the pacemaker model cannot explain a large body of observations that have accumulated over the past decades. First, hippocampal neurons are not locked synchronously to a global rhythm but show a systematic phase shift up to 270° in the CA1-CA3-dentate gyrus axis and in different layers of the entorhinal cortex (Buzsáki et al., 1986; Mizuseki et al., 2009). Second, theta is not synchronous over the entire septotemporal axis but, instead, shows a traveling wave pattern, spanning 180° from the septal to the temporal pole and (Lubenov and Siapas, 2009; Patel et al., 2012). Third, all activated principal cells, such as neurons that fire at particular spatial positions ‘place cells’ (O’Keefe and Nadel, 1978; O’Keefe and Recce, 1993) or in a given memory episode (Pastalkova et al., 2008), oscillate faster than the LFP theta. The oscillation frequency of place cells correlates inversely with the diverse sizes of place fields (Dragoi and Buzsáki, 2006) and vary systematically along the septotemporal axis (Kjelstrup et al., 2008; Maurer et al., 2006; Royer et al., 2010). Similarly, theta oscillation frequency of ‘grid cells’ in the entorhinal cortex decreases progressively in the dorsoventral direction (Giocomo et al., 2007), providing a frequency match between corresponding entorhinal and hippocampal neurons. Finally, as a result of the phase interference between population spiking behavior, as reflected by LFP theta, and the faster oscillatory spiking of individual neurons (Skaggs et al., 1996; Geisler et al., 2010) show a progressive backward phase shift of their spikes as their activity unfolds (‘phase precession’) (O’Keefe and Recce, 1993). In summary, spikes of principal cells in the limbic system occur at all phases of the theta cycle and all active principal cells oscillate faster than the global LFP theta whose instantaneous frequency co-varies coherently across subregions and structures (Figure 1). These experiments suggest a more elaborate involvement of MS circuit in theta cycle phasing of hippocampal neuronal assemblies than the current models would imply (Borhegyi, 2004; Buzsáki, 2002; Hangya et al., 2009; Stewart and Fox, 1990).

**Fig. 1.**
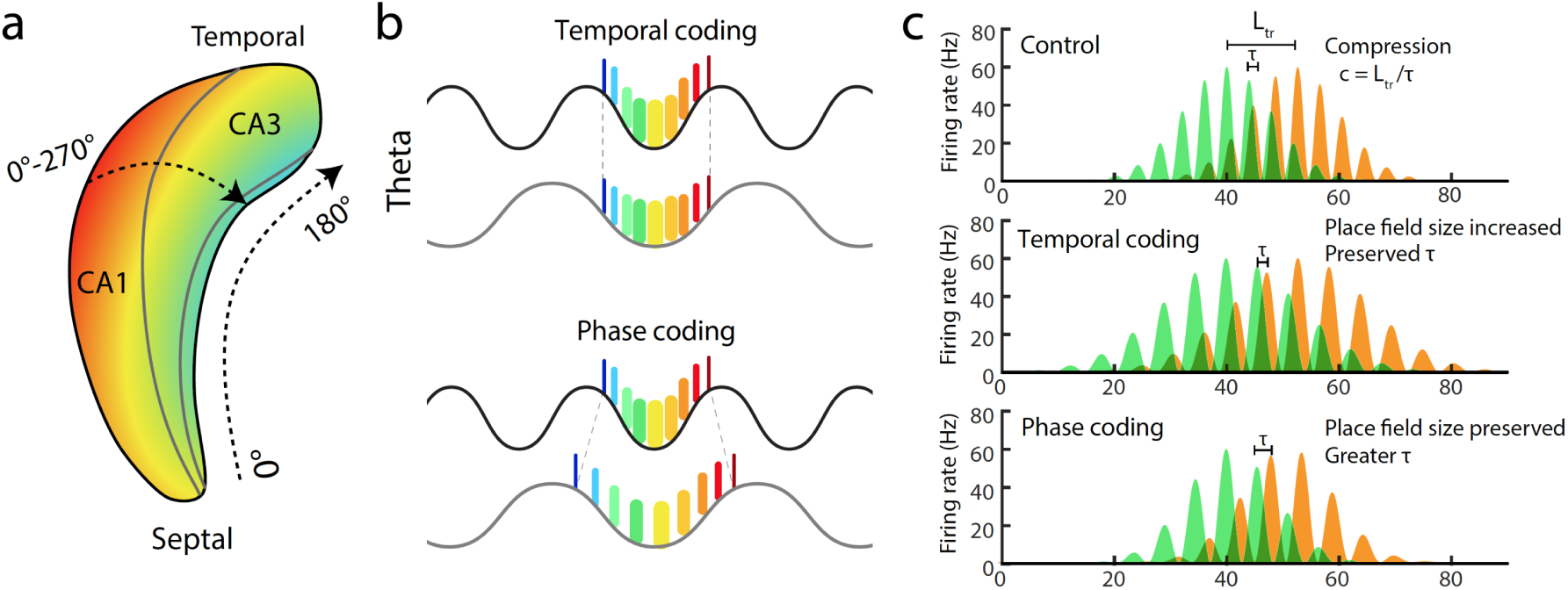
Theta phase dynamics and connectivity in the MS-hippocampal system. **a**, Theta oscillation is a traveling wave and shows systematic phase shift in both the septo-temporal and CA1–CA3–dentate gyrus axes (color scale illustrates the phase offsets between regions). Hippocampal neurons are not locked synchronously to a global theta rhythm but show a systematic phase shift up to 270° in the CA1-CA3-dentate gyrus axis and a gradual 180° shift from the septal to the temporal pole (Lubenov and Siapas, 2009; Patel et al., 2012). **b**, Hypothetical effects of MS cooling on within-theta neuronal assembly organization. Top: Time lags between place cells assemblies (different color ticks) remain unchanged with theta cycle lengthening (gray). Bottom: Place cells assemblies expand in time but keep their theta phase relationships. **c**, Theta cycle relationship between two overlapping place fields of the same size (*L*_*PF*_). The travel distance (*L*_*tr*_) between the peaks of the place fields is correlated with the time offset (τ) of spikes between the green and tangerine place neurons (Dragoi and Buzsáki, 2006). The ratio of travel distance and theta time scale time lag defines distance to time compression (*c* = *L*_*tr*_ / τ) (Geisler et al., 2010). Middle and bottom, Hypothetical effects of MS cooling. Middle: Place field size changes but τ is preserved. Bottom: Fewer theta cycles occur within the same size place field and τ increases.

Damaging, silencing or pharmacological perturbation of MS circuit (Bolding et al., 2020) cannot effectively address temporal/theta phase coordination issues. Perturbation studies using synchronizing electrical (McNaughton et al., 2006) or optogenetic (Dannenberg et al., 2019; Vandecasteele et al., 2014; Zutshi et al., 2018) stimulation may not be effective either to fully examine this problem because strong pulses impose global synchrony on all neurons unlike the time-shifted patterns observed under physiological conditions (Lubenov and Siapas, 2009; Patel et al., 2012). Therefore, we used temperature manipulation, an approach that is applicable to localizing the origin of temporal coordination (Fee and Long, 2011). In contrast to unwanted synchronizing stimulation, cooling does not interrupt local interactions but alters multiple parameters of neurons from channel kinetics to transmitter release, resulting in temporal warping of circuit dynamic (Katz et al., 2004; Thompson et al., 1985; Volgushev et al., 2000). For example, reducing temperature in the vocal center of the zebra finch elongated the bird’s song by proportionally slowing its acoustic microstructure (Long and Fee, 2008).

By cooling the MS, we examined how slowing theta oscillations affect hippocampal network activity, physiological properties of neurons and their spatial correlates (O’Keefe and Nadel, 1978). Several models assume that the phase interference between MS theta oscillation (LFP theta) and place inputs-driven faster oscillations of place cells and grid cells determine the slope of spike phase precession, and, consequently, place field size (Burgess et al., 2007; Chadwick et al., 2016; Harvey et al., 2009; Kamondi et al., 1998; O’Keefe and Recce, 1993; Zutshi et al., 2018). A prediction of these models, therefore, is that altering MS-driven theta oscillation should affect the size of place fields. Furthermore, if the temporal lags within cell assemblies are preserved, more assemblies can be packaged in a wider theta cycle (‘temporal coding’). An alternative hypothesis is that the fundamental organization in the septo-hippocampal system is not time-but phase-based (‘phase coding’). Under the phase-model, theta phase assembly coordination should remain unaltered but at the expense of affecting timing between place cell assemblies (Figure 1). Our results favor the phase coding model of theta cycle coordination.

## Results

### MS cooling decreases hippocampal theta frequency and power

To achieve localized cooling of the medial septum, we constructed a cryoprobe, consisting of a silver wire (125µm in diameter), 25µm graphene sheet, air isolation and a polyimide tube (Fig. 2a; see Methods). The back end of the silver wire was coiled at the bottom of an insulated reservoir. Cooling was achieved by filling the reservoir with dry ice (−78.5° C). The protruding tip (1.5 mm) of the front-end wire was implanted into the medial septum-diagonal band area (MS; Fig. 2b). A temperature sensor (k-type thermocouple; two 80µm wires) was attached to the outer surface of the cryoprobe cannula to continuously monitor local temperature (Fig. 2a; Suppl. Fig. 1). Pilot experiments were performed to perfect the device so that hippocampal theta frequency could be reduced for 5 to 15 min (Suppl. Fig. 1).

**Fig. 2.**
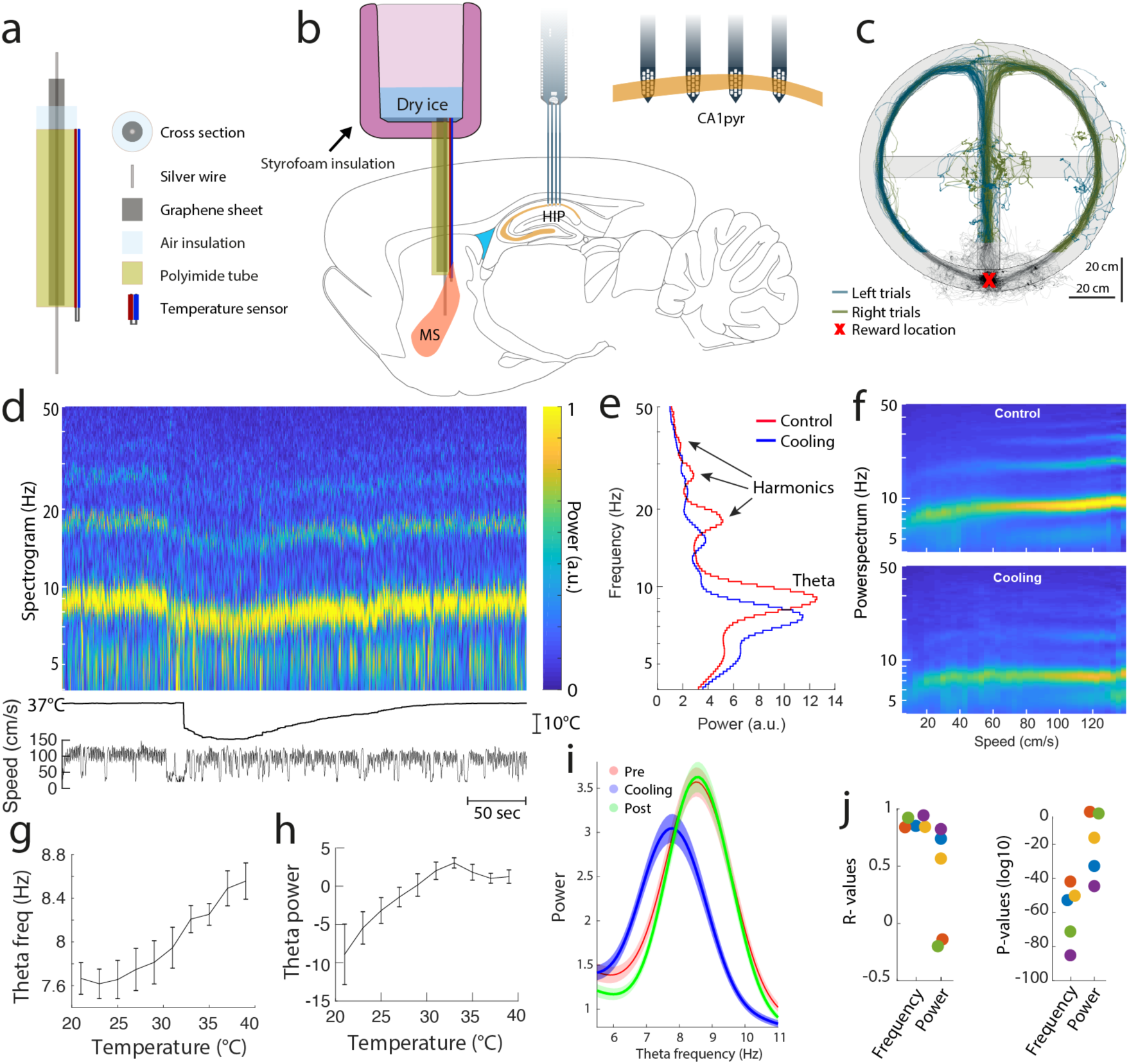
Cooling of MS affects theta oscillations. **a**, Components of the cooling probe. **b**, Cooling probe and thermocouple are implanted in MS and silicon probes in the hippocampus. Dry ice is placed into the reservoir and MS is cooled by thermal conduction. **c**, Maze with left (blue) and right (green) run trajectories superimposed. **d**, top panel: Time-resolved spectrogram in a single session. Color scale is normalized. Middle panel: temperature; lower panel: running speed of animal. **e**, Frequency spectrum with power of theta and its 1^st^-3^rd^ harmonics (arrows). **f**, Power spectra of the theta band and harmonics before (top) and after (bottom) cooling. Color scale is normalized. Same single session used in d,e,f. **g, h**, Effect of MS cooling on theta frequency (**g**) and power (**h)** Group data for all sessions and subjects (mean values with SE)**. i**, Group data for theta frequency and power before, during and after MS cooling. **g**,**h**,**i**,**j**: 53 sessions in 5 rats. **j**, R and P values for theta frequency and power in individual rats (color coded).

In addition to the cooling probe, five rats were also implanted with multi-shank 64-site silicon probes bilaterally in the CA1 pyramidal layer of the dorsal hippocampus (Supplementary Material) and were trained in a figure-8 maze (spontaneous alternation task; Fig. 2b) to run for water reward. After a block of control trials (∼ 40 trials), dry ice was placed into the reservoir, which reduced MS temperature for approximately 15 min (or corresponding to about 20 to 60 ‘cooling’ trials), followed by recovery trials (20 to 80). MS cooling induced fast and reversible change in the frequency and power of hippocampal theta oscillations (Figure 2d, e, f, i). Theta frequency decreased linearly with MS temperature (Figure 2g; down to 20° C; R = 0.81; P = 5.4 × 10^−48^) and this relationship was more consistent across animals than the effect on theta power (R = 0.54; P = 1.2 × 10^−16^; Fig. 2g, h). The time course of theta frequency decrease, and less so its power, mirrored the temporal dynamic of MS cooling (see time course in Fig. 6).

**Fig. 3.**
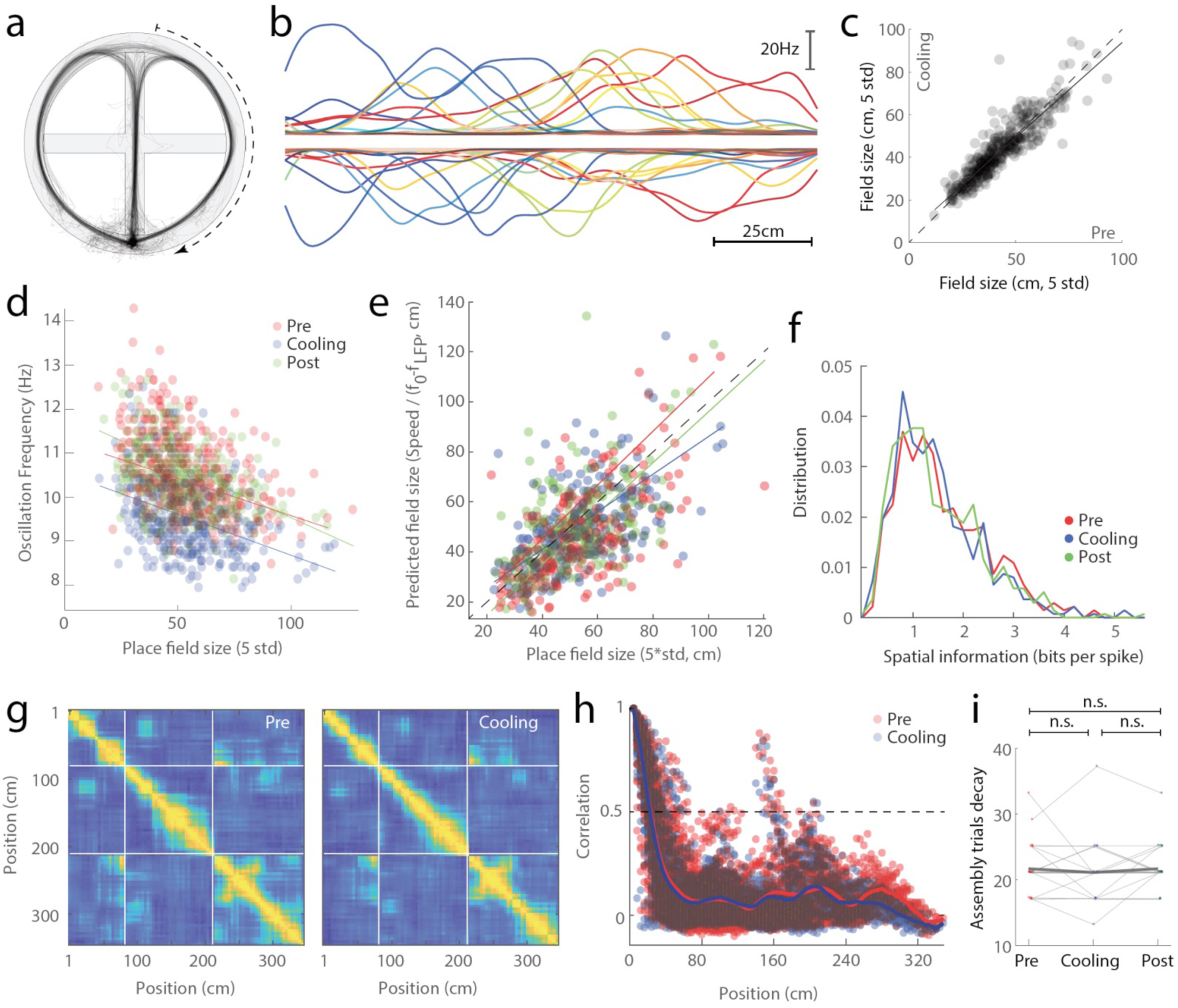
Place field features are not affected by MS cooling. **a**, Example of right arm runs in the T maze. **b**, Place fields in the right arm before (upward traces) and during MS cooling (downward traces) in a single session. Note preserved size, position and shape of place fields. **c**, Place field size does not change with MS cooling (R = 0.91). **d**, Place field size continues to be inversely correlated with f_p_ after MS cooling (R_pre_ = -0.49; R_cooling =_ - 0.40; R_post_ = 0.40. P < 10^−9^ all conditions). **e**, Correlation between measured (x) and model-predicted (y) size of place fields before, during and after MS cooling. Place field size in the model (Geisler et al., 2010) was derived from *S*_*PF*_ = *v*/(*f*_*p*_ – *f*_*LFP*_). *v* = speed. **f**, Information content of place cell spikes. None of the comparisons are significantly different (P > 0.26). **g**, Population vector cross-correlation matrices from a single session from baseline (pre-cooling) and MS cooling trials. White lines mark the boundary between central arm, right and left arms. Normalized color scales are the same in the two panels. **h**) Superimposed decorrelation curves for trials before and during MS cooling. To quantify the scale of the spatial representation, the interval at which the correlation dropped to 0.5 was calculated for each session (dashed line). **i**) Decorrelation values for data from 21 sessions.

**Figure 4.**
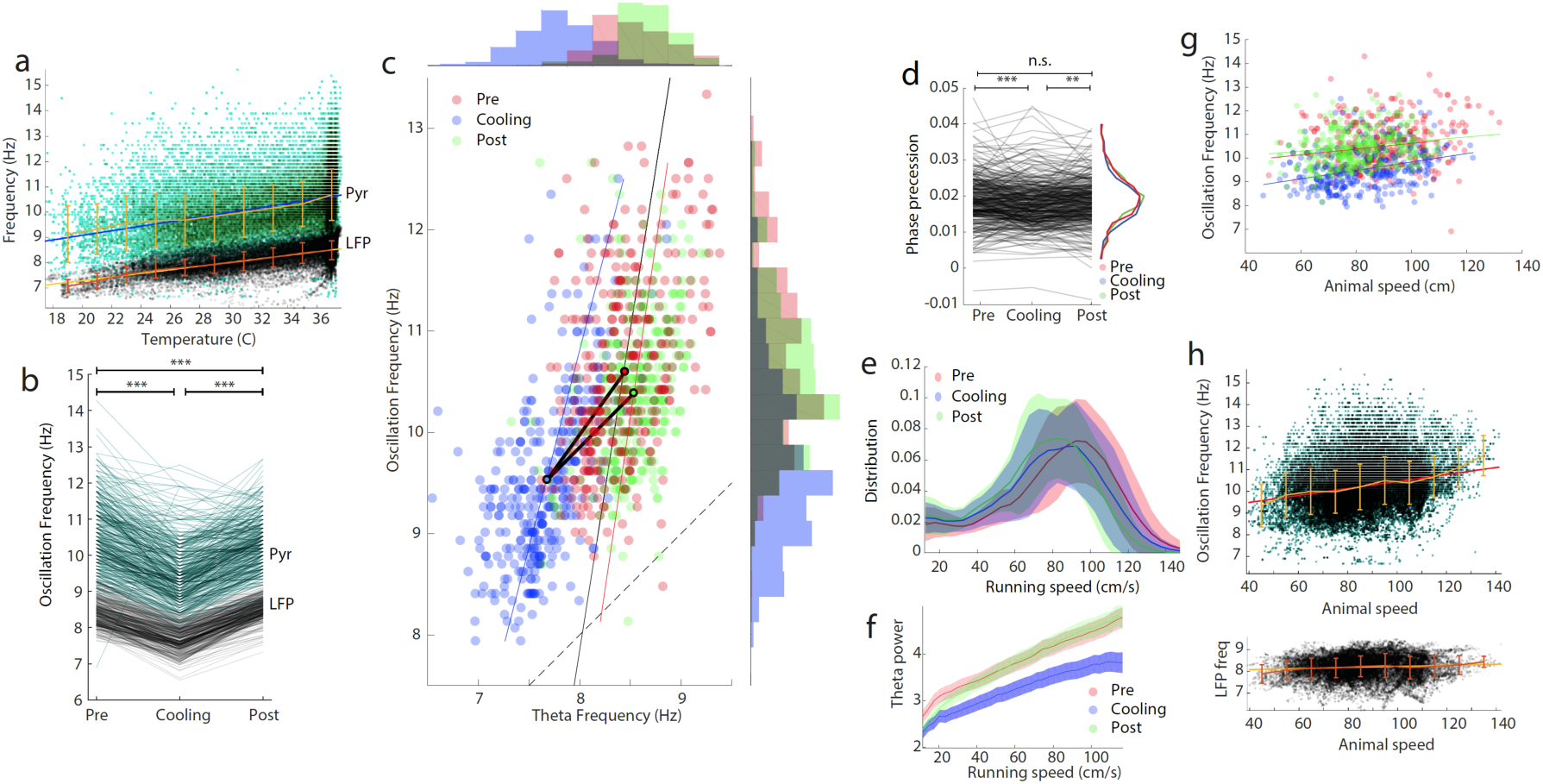
Effects of MS cooling on single unit parameters. **a**, LFP theta frequency (*f*_LFP_) and oscillation frequency of place cells (*f*_*p*_) as a function of MS temperature. Each dot is a trial average measured within a single place field. Note parallel changes between *f*_*p*_ (green) and *f*_*LFP*_ (black). **b**, Oscillation frequency shown separately for all place cells (*f*_*p*_; gray) and LFP theta frequency (*f*_*LFP*_; black) measured in the corresponding place fields before, during and after MS cooling. *** P < 5.9e–8 **c**, Correlation between LFP theta frequency (*f*_*LFP*_) and place cell oscillation frequency (*f*_*p*_; R_pre_ = 0.61; R_cooling_ = 0.59; R_post_ = 0.36; P < 10^−19^ each). Black lines connect the centers of mass of each cloud. Marginal histograms show the distribution of *f*_*p*_ of place cells and *f*_*LFP*_ in the corresponding place fields. Dashed line: diagonal. **d**, Theta phase precession slopes of place cell spikes before, during and after MS cooling. ** P < 0.007; *** P < 0.0003. **e**, Distribution of running speeds in all sessions (N = 53). **f**, Theta power as a function of running speed. **g**, Place cell oscillation frequency (*f*_*p*_) as a function of running speed, measured in each place field. **h**, *f*_*p*_ and *f*_*LFP*_ as a function of running speed. As in **g** but trials before, during and after MS cooling are pooled to show that *f*_*p*_ depends more strongly on running speed than *f*_*LFP*_.

**Figure 5.**
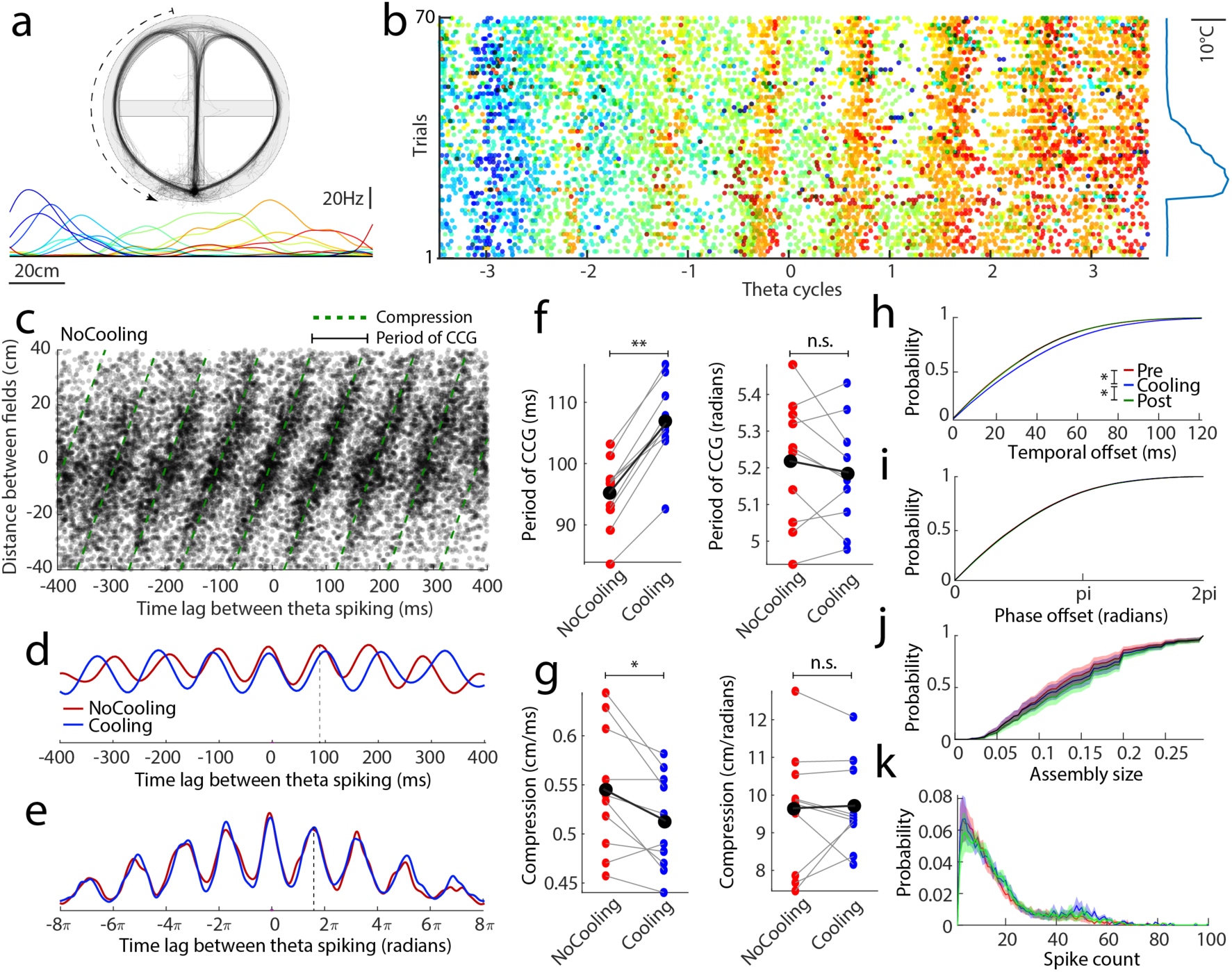
Distance–theta time scale compression is affected by MS cooling. **a**, Selected place fields in the left arm of the maze before MS cooling in a single session. **b**, Within-theta cell assemblies. Each row of dots is a trial of the spiking activity of place cells (same color code as in **a** in successive theta cycles). 0 is the theta cycle in the middle of the left arm. Note shifting cell assemblies in successive theta cycles (from blue to red) and preserved phase preference of place cell spikes during and after MS cooling (right trace). **c**, Relationship between distances of place field peaks across neuron pairs (y axis) and their theta time-scale cross-correlogram lags (τ on x axis; as in Fig. 1c) during trials prior to MS cooling (2948 = pairs across 18 sessions, 5 rats). Green dashed line; compression index (slope), *c =* 0.54 cm/ms. Period of the CCG highlighted (black solid line). **d**, Sum of the theta repeated spike cross-correlograms of place field pairs. The red line is the sum of all dots in **c** (NoCooling); blue line, sum of all dots of a similar graph during cooling. Vertical dashed line indicates that the oscillation frequency of the spike cross-correlation before cooling (∼ 85 ms) is faster that *f*_*LFP*_ (∼110 ms at 9 Hz). **e**, Similar to **d** but instead of time lags, within-theta phase lags between spikes were calculated. Vertical dashed line indicates that the oscillation frequency of the spike cross-correlation is faster than that of *f*_*LFP*_ (2π). **f**, The duration of the theta-repeated spike cross-correlograms (as in **d**), calculated for each session, significantly increased during MS cooling, compared to control (Pre and Post-cooling epochs combined) trials (** P = 0.002; two-sided signed rank test), whereas the spike phase-lag correlations remained unaffected (n.s., P = 0.32). Black disks and connected line, group means. **g**, Compression index, *c* (cm/ms; i.e., the slope in **c**), significantly decreased during MS cooling (* P = 0.02) but the distance/phase lag compression (cm/radian) remained unaffected (P = 0.92). **h**, Cumulative distributions of theta scale time lags (τ) between place cell pairs before (Pre), during and after (Post) MS cooling. MS cooling data are significantly different from both Pre- and Post-cooling trials (Pre vs cooling P = 8.0^−5^; Post vs cooling P = 0.00032; N = 10 sessions, 4 rats). Pre and post-cooling curves are strongly superimposed (Pre vs Post P = 0.79). **i**, Same as in **h**, but for phase lags (all comparisons are non-significant, P > 0.24). **j**, Cumulative plots of the fraction of all simultaneously recorded neurons active in the theta cycle (pre vs cooling P = 0.28; post vs cooling P = 0.006; pre vs post P = 0.0005). **k**, Distribution of spike counts within theta cycles (Pre and Post-cooling epochs are not different from MS cooling epochs or from each other; P > 0.058). y axis, probability.

**Figure 6.**
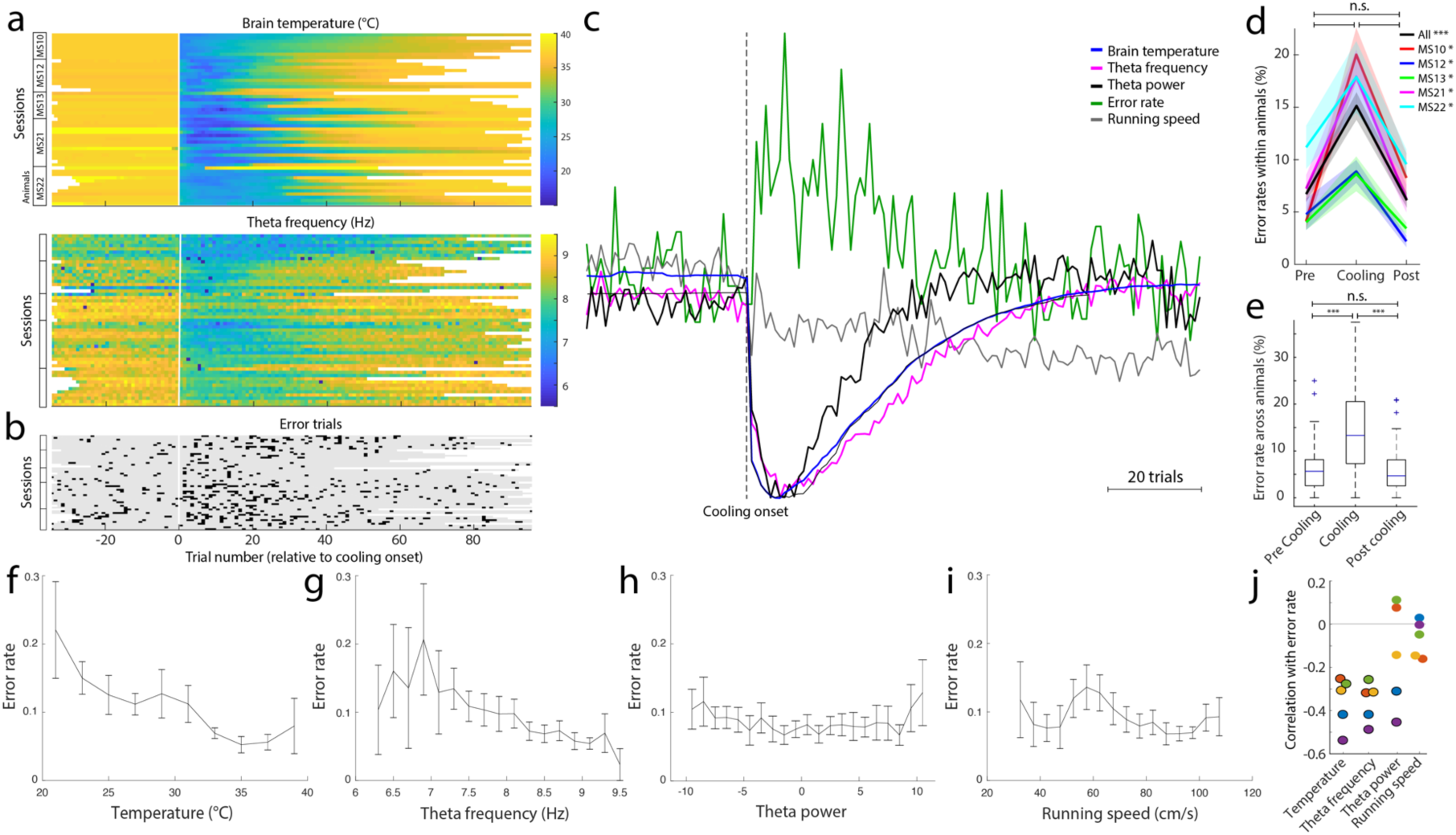
Effects of MS cooling on memory performance. **a**, MS temperature (top) and theta frequency shift (bottom) as a function of trial number (x axis). Each line corresponds to a single session. Animal IDs are indicated on the left. Vertical line marks the onset of cooling. **b**, Incidence of error trials in the same sessions. **c**, Time courses of MS temperature, theta frequency, power and running speed (mean of all sessions). Note tight correlation between MS temperature, theta frequency and power. In contrast, speed shows a monotonic function across trials. **d**, Choice errors shown separately for each animal. Standard errors are shown in shaded colors. Black line, group means. * P < 0.05; *** P = 10^−6^. **e**, Choice errors shown as boxplots constructed from all sessions. ** P < 10^− 7^. ns = nonsignificant. **f-i**, Relationship between memory errors and temperature (**f**), theta frequency (**g**), theta power (**h**) and running speed (**i**). **j**, Correlation values between memory errors and parameters shown in **f** to **i** for each rat. Significant values indicated by black circles (P < 0.05). Same color indicates same animal.

In a control experiment, we measured temperature simultaneously in the MS and hippocampus, and found < 1.5° C decrease in the hippocampus at the time of the largest MS temperature decrease (16° C), that is within the physiological range of brain temperature (Suppl. Fig. 1h-k) (Moser and Andersen, 1994). Critically, in contrast to the rapid onset of cooling in MS (maximum effect at approximately 45 s; time constant: 11 s), the mild temperature decrease in the hippocampus was considerably delayed (peak at 115 s; time constant 48s for the right hippocampus, and 72 s for the left). Furthermore, firing rates of both pyramidal cells and putative interneurons in the hippocampus decreased immediately after MS cooling and the changes correlated well with MS temperature change (see Suppl. Fig. 5g, h) rather than with the delayed minimal temperature decrease in the hippocampus. Finally, the waveforms of action potentials in the hippocampus, a sensitive measure of temperature change (Thompson et al., 1985; Volgushev et al., 2000), were not affected by MS cooling (Suppl. Fig. 1g). Overall, these findings show that MS cooling had an immediate and strong effect on LFP theta and suggest that these changes were not a consequence of the spread of cooling to the hippocampus.

### Effect of MS cooling on place field features

Spatial parameters of place fields were largely left unaffected by MS cooling. Place field size (*L*_*PF*_) remained highly correlated before and after MS cooling (Fig. 3a, b; see Suppl. Fig. 3 for other place field features). *L*_*PF*_ and *f*_*p*_ were negatively correlated (Dragoi and Buzsáki, 2006; O’Keefe and Recce, 1993), and this relationship was not altered by cooling, as illustrated by the similar slopes, even though the intercept of the regression line during cooling slightly decreased (Fig. 3c, d).

Next, we compared measured place field size with model-based calculations. Previous studies have demonstrated a quantitative relationship among theta frequency (*f*_*LFP*_), oscillation frequency of pyramidal cells within their place fields (*f*_*p*_), the duration it takes the rat to run across the place field (*D*_*PF*_), place field size (*S*_*PF*_) and the rat’s velocity (*v*) (Geisler et al., 2010). Since *D*_*PF*_ = *1/f*_*p*_ – *f*_*LFP*_ (Geisler et al., 2010) and *D*_*PF*_ can be calculated by dividing place field size (*S*_*PF*_) with locomotion velocity within the place field (*S*_*PF*_ = *D*_*PF\**_*v*), place field size can be derived from *S*_*PF*_ = *v*/(*f*_*p*_ – *f*_*LFP*_). We found a reliable and similar relationships between measured place field sizes and model-predicted sizes for trials before (R = 0.55), during (R = 0.54) and after (R = 0.11) MS cooling (Fig. 3e). Information content of place cell spikes was also left unaltered by MS cooling (P > 0.26; Fig. 3f).

We also investigated potential changes to the scale of distance representation at the population level resulting from MS cooling. We calculated the correlations between pairs of population vectors of unit firing constructed from binned rate maps (Gothard et al., 1996). The distance in maze corridor bins at which the spatial population vector correlation function dropped to 0.5 was used as a measure of the scale of the spatial representation, a measure independent of arbitrary definition of place field (Gothard et al., 1996). Population vector correlation of pyramidal cells was not affected by MS cooling (Fig. 3g, h). This analysis further suggested that spatial representation by place cell population was largely unaffected by MS cooling (Fig. 3i).

### Effects of MS cooling on single neuron properties

In contrast to the lack of changes of spatial features, MS cooling induced widespread but correlated physiological effects (Fig. 4). The theta oscillation frequency of pyramidal neurons decreased linearly and paralleled the downward frequency shift of LFP theta oscillation (Fig. 4a; *f*_*LFP*_: R=0.66, Slope=0.071. *f*_*p*_: R=0.38, slope=0.09). Theta frequency oscillation of putative fast spiking interneurons also decelerated (Suppl. Fig. 2d), commensurate with the downward frequency shift of LFP theta oscillation.

The range of within-field oscillation frequency of place cells (*f*_*p*_) was larger compared to the range of LFP theta frequency within place fields (*f*_*LFP*_; Fig. 4c), indicating altered theta cycle representation of place fields. MS cooling decreased both *f*_*p*_ and *f*_*LFP*_, measured within the place fields of neurons, by approximately the same degree (Fig. 3b,c; *f*_*LFP* pre_=8.4±0.38, *f*_*LFP* cooling_=7.7±0.36, *f*_*LFP* post_=8.5±0.29; *f*_*p* pre_=10.6±0.95, *f*_*p* cooling_=9.5±0.84, *f*_*p* post_=10.4±0.72, n=365). Both values recovered after cooling. The *y* intercepts of regression lines of the *f*_*p*_ versus *f*_*LFP*_ relationship shifted from pre-cooling to cooling but remained parallel to each other in each condition. The lines connecting the centers of masses also remained largely parallel with the diagonal (Fig. 3c), indicating preserved *f*_*p*_ – *f*_*LFP*_ differences. Both within-field and peak firing rates of place cells decreased moderately during MS cooling (Suppl. Fig. 2; 11% and 12%; P < 10^−9^ and P < 10^−11^, respectively; see time course in Suppl. Fig. 6). Interneuronal firing rates also decreased by approximately the same degree (Suppl. Fig. 2). The slope of the relationship between travel distance across the place field and phase advancement of spikes (‘phase precession’ index) (O’Keefe and Recce, 1993) decreased slightly, but nevertheless significantly, during MS cooling (Fig. 4d).

Since previous studies showed that the running speed of the animal is correlated with several physiological parameters (Dannenberg et al., 2019; Diba and Buzsáki, 2008; Fuhrmann et al., 2015; Geisler et al., 2007; Maurer et al., 2012; McNaughton et al., 1983), we aimed to disambiguate the effect of MS cooling and running speed. Running speed decreased relatively monotonically within session (Fig. 4e; see also Fig. 6c). The animal’s speed correlated with both theta power and *f*_*p*_ (Fig. 4f, g). Yet, *f*_*p*_ changes during MS cooling could not be explained by decreased running velocity of the rat because *f*_*p*_ values during cooling were lower at all velocities (Fig. 4g). We examined the relationship between running speed and LFP theta frequency (*f*_*LFP*_) in two different ways. The first comparison was done on a trial basis, comparing the mean theta frequency with the mean running speed in each trial. This comparison showed a positive correlation before, during and after MS cooling with the largest changes taking place at lower running speeds (10 to 40 cm/s; Suppl. Fig. 2e). However, this comparison cannot dissociate the effect of speed from the effect of maze environment (Montgomery et al., 2009). Therefore, we quantified speed – theta frequency relationship within each place field (*f*_*LFP*_), together with the oscillation frequency of the corresponding place cell (*f*_*p*_). This calculation revealed that while speed exerted a relatively linear effect on *f*_*p*_, it had no or little effect on the frequency of *f*_*LFP*_ (Fig. 4h) (Czurkó et al., 1999; Montgomery et al., 2009) within the dominant speed range of the rat. As a result, the *f*_*p*_ – *f*_*LFP*_ difference increased with running velocity. This is in contrast to the effect of MS cooling, after which the difference between *f*_*p*_ and *f*_*LFP*_ was preserved (Fig. 4a and c). Finally, we examined duration of theta cycles as function of the rat’s position on the track. While theta duration was consistently longer during MS cooling trials, duration distribution did not correlate with the speed distribution on the track (Suppl. Fig. 3). Overall, these findings indicate that the physiological effects of MS cooling are dissociable from running speed effects (Tsanov, 2017).

### MS temperature effects on theta time/phase compression of place cells sequences

Our findings so far revealed a contrast between consistent changes in physiological parameters of single neurons and the stability of their place field features by MS cooling. Because single neuron features do not inform us about their theta-organized assembly properties, we next examined the temporal sequences of place cell assemblies in the theta cycle (Fig. 5a, b). The distances between place cell peaks (*L*_*tr*_) are known to correlate with their theta-scale time lags (τ) and phase lags, and the ratio of *L*_*tr*_ and τ is known as distance-time compression (*c*; figure 1a) (Dragoi and Buzsáki, 2006; O’Keefe and Recce, 1993). To display the dynamics of ensemble compression of place cell distances as the animal passes through sequential positions on the maze, we plotted the sequence compression for the entire population of place cell pairs over several theta cycles within a 0.8-s period. In Fig. 5c, each dot corresponds to the time difference of spike occurrences of place neuron pairs (τ), averaged across all trials of a session (n = 18 sessions in 5 rats). This representation mimics the spatial-temporal evolution of spiking of all pyramidal neurons in the dorsal hippocampus as the rat traverses the place field center of a reference neuron (Dragoi and Buzsáki, 2006). The “clouds” are spaced by theta-scale intervals, relating to the joined oscillating frequency of place cell pairs, and the slope of the clouds corresponds to the distance-to-theta scale-time compression (*c*) (Maurer et al., 2012). Several place cells are active together in each theta cycle, but the group composition varies from cycle to cycle (Fig. 5b) (Dragoi and Buzsáki, 2006; Maurer et al., 2012)

Both the joined oscillating frequency of place cell pairs and *c* were affected by MS cooling (Fig. 5d, f, g). Because the compression index also depends on running speed (Maurer et al., 2012) and because running speed gradually decreased across the entire session (Fig. 3e; 6c), we combined trials before and after cooling into a single control group (mean running speed = 87.3±8.0). During MS cooling (mean running speed = 86.2±9.9), 1/*f*_*p*_ increased (i.e., *f*_*p*_ decreased) in each examined session relative to control runs (Fig. 5f, g; 0.002; Wilcoxon signed rank test), whereas distance-time compression decreased (Fig. 5g; P = 0.02). We also constructed analogous plots displaying *L*_*tr*_ versus theta phase lags of spike pairs, instead of time lags. Phase lags remained unchanged after MS cooling (Fig. 5f; P > 0.24 for all pairs; Wilcoxon signed rank test) and, as a consequence, the ratio (i.e., *L*_*tr*_ versus theta ‘phase compression’) between *L*_*tr*_ and phase lag did not change either (P < 0.92; Fig. 5g).

Because distances between place field peaks (*L*_*tr*_) were not affected by MS cooling (Fig. 3c and g-i) but distance-time compression decreased (Fig. 5g), time lags (τ) are expected to increase. To examine this effect directly, we plotted the distributions of both theta time lags (τ) and phase lags between all pairs of place cells in each session. During MS cooling, the time lags increased significantly (Fig. 5h; P < 10^−4^; sign rank test; n = 10 sessions), whereas they were not different from each other during precooling and post-cooling sessions (P = 0.79), despite slower running speed during Post compared to Pre-cooling trials. In contrast, phase lags between spikes of place cell pairs were not affected by MS cooling (Fig. 5i; P > 0.24 between all conditions). Thus, theta phase lags remained similar while time lags increased.

These findings suggested that the same number of place cell assemblies were compressed into theta cycles both before after MS cooling and the assemblies were proportionally dispersed within the lengthened cycle during cooling trials (Fig. 5c,d). In support of this hypothesis, we found that the fraction of place cells active in a given theta cycle was not significantly affected by MS cooling (Fig. 5j; P = 0.2). The number of spikes emitted by all neurons per theta cycle was also preserved after MS cooling (Fig. 5k), which might be explained by the similar magnitude of the within-field firing rate decrease of place cells (∼12%; Suppl. Fig. 2) and theta frequency decrease (∼12%; Fig. 4a and b; trial-wise correlations, see Suppl. Fig. 5).

### MS cooling increases behavioral errors

The temperature decrease in the MS was immediately accompanied by increased choice errors in the maze (Fig. 6a-c). The worst memory performance occurred between trials 4 and 8 after MS cooling, when the MS temperature reached its minimum, accompanied by the largest decreases of theta frequency and power (Fig. 6c; see also Suppl. Fig. 5). Overall, the number of memory errors increased approximately 3-fold, assessed either within rats (P < 0.05 for all animals) or for all animals (P = 10^−6^; Fig. 6d, e). The incidence of errors negatively and significantly correlated with temperature change (R = – 0.35; P < 0.01; all animals), theta frequency (R = – 0.35; P < 0.01; all animals) and less so with theta power (only 2 of 5 animals had significant R > – 0.3; P < 0.01) and not with the running speed of the rat (R ∼ 0.1; P > 0.05 for all animals; Figure 6f-j; see relationship to firing rates and number of spikes per theta cycle in Supp. Fig. 5).

## Discussion

Cooling the MS reduced both LFP theta frequency and power in the hippocampus, associated with enhanced incidence of errors in a spatial navigation task without affecting spatial attributes of individual place cells and the spatial map. The slowing of LFP theta oscillation was paralleled with a commensurate a) decrease of theta frequency oscillation of place cells, b) theta time cross-correlations of place cell pairs, c) reduced time compression but preserved phase compression of place field sequences within theta cycles and e) a decrease of firing rates of both pyramidal cells and interneurons, commensurate with the increased duration of theta cycles.

MS cooling may non-specifically affect several targets, including local neurons and their intraseptal interactions, transmitter release probability subcortical neuromodulators terminating in MS and it can also alter spike propagation in axons passing through the MS (Fee and Long, 2011; Volgushev et al., 2000). Our goal was not to disclose the local physiological effects of temperature manipulations in MS but to bring about a reliable macroscopic phenotype, theta frequency, and examine the circuit mechanisms responsible for such change in the hippocampus. Since the cooling method affects circuit components relatively homogeneously, it is an alternative or complementary tool to neuron-specific perturbations (Fee and Long, 2011; Fuhrmann et al., 2015; Lee et al., 1994; Vandecasteele et al., 2014; Zutshi et al., 2018). Many of the subcortical neuromodulatory and supramammillary effects are likely mediated by affecting MS neurons (Vertes and Kocsis, 1997). They may be responsible for altering theta power but less so theta frequency. Selective damage to MS cholinergic neurons reduces theta power several-fold without affecting theta frequency (Lee et al., 1994). Furthermore, pharmacological blockade of muscarinic cholinergic receptors does not alter the cross-correlation structure of neuron pairs, spatial properties of place cells, the predictability of spiking in individual place cells from peer activity, or the decodability of patterns of simultaneously recoded place cell spikes (Venditto et al., 2019). Conversely, optogenetic activation of MS GABAergic, but not cholinergic, neurons can tune theta frequency (Zutshi et al., 2018; Fuhrmann et al., 2015), suggesting that MS cooling exerted its theta temporal/phase effects mainly via affecting MS GABAergic neurons (Borhegyi, 2004; Hangya et al., 2009; Simon et al., 2006; Stewart and Fox, 1990; Zutshi et al., 2018). Thus, even if MS cooling affects multiple mechanisms, the strongest correlation between MS temperature and theta frequency oscillation suggests that MS GABAergic neurons are likely involved.

### Theta phase organization of hippocampal cell assemblies

Early experiments suggested that the MS acts as a ‘pacemaker’, sending out synchronous outputs, akin to a conductor of an orchestra (Borhegyi, 2004; Buzsáki, 2002; Chadwick et al., 2016; Petsche et al., 1962; Stewart and Fox, 1990; Sweeney et al., 1992; Zutshi et al., 2018). An alternative model is based on the reciprocal relationship between MS and the hippocampus (Hangya et al., 2009). In one arm of the loop, MS GABAergic neurons innervate hippocampal and entorhinal interneurons (Freund and Antal, 1988; Fuchs et al., 2016; Gonzalez-Sulser et al., 2014; Takács et al., 2008; Unal et al., 2015). In the return direction, long-range GABAergic neurons send axons to the MS (Gulyás et al., 2003; Toth et al., 1993).

Interneurons, populations of pyramidal cells and MS neurons all fire at the frequency of hippocampal LFP (Buzsáki, 2010; Geisler et al., 2010; Gothard et al., 1996). In contrast, active individual pyramidal cells and granule cells oscillate faster (*f*_*p*_) than the population activity (*f*_*LFP*_), resulting in phase interference and measured experimentally by the ‘phase precession’ slope (O’Keefe and Recce, 1993). The difference between oscillation frequency of place cells and LFP theta is inversely correlated with place field size and affected by the running speed of the animal (*L*_*PF*_ = *v*/(*f*_*p*_ – *f*_*LFP*_); (Geisler et al., 2010). We found that at faster velocities the difference between *f*_*p*_ and *f*_*LFP*_ increases, which may explain the place field size invariance (Geisler et al., 2007; Huxter et al., 2003).

If the sole function of MS were to supply a pacemaker pattern by entraining target interneurons in the hippocampus (Freund and Antal, 1988) at LFP frequency and if LFP frequency and oscillation frequency of place cells were independent (as assumed in (Chadwick et al., 2016), then slowing the output MS theta frequency would result in a larger *f*_*p*_ – *f*_*LFP*_ difference and, consequently steeper phase precession and smaller place fields. However, during MS cooling *f*_*p*_ and *f*_*LFP*_ decreased proportionally (i.e., *f*_*p*_ – *f*_*LFP*_ difference remained constant) and the influence of the rat’s running velocity on *f*_*p*_ was preserved (Fig. 4a, c). These preserved relationships can explain why place field size was not affected by MS cooling. Proportional slowing of *f*_*p*_ and *f*_*LFP*_ was also observed in a virtual reality experiment, suggesting that theta phase preservation of spikes is a fundamental mechanism (Aghajan et al., 2015; Ravassard et al., 2013).

The travel distances between the peak positions of place cell sequences (*L*_*tr*_) are correlated with their theta-scale time lags (τ) and phase lags, and the ratio of travel distance and theta-scale time lag (*L*_*tr*_/τ) is known as distance-to-time compression (*c)* (Dragoi and Buzsáki, 2006; Maurer et al., 2012). This distance-to-duration compression is believed to be the mechanism that chains together evolving cell assemblies, and explains the succession of place fields during navigation emerging from place cell sequences within theta cycles (Chadwick et al., 2016; Dragoi and Buzsáki, 2006; Harris et al., 2002; Lisman and Jensen, 2013; Mehta et al., 2002; O’Keefe and Burgess, 2005; Pastalkova et al., 2008; Samsonovich and McNaughton, 1997; Skaggs et al., 1996; Wang et al., 2015). Place field sizes (*L*_*PF*_) and the distances between place field peaks (*L*_*tr*_) were not affected by MS cooling. On the other hand, the theta-scale time lags (τ) between successive place cell assemblies increased. As a result, the compression index (*c*) decreased. In contrast, phase lags between successive place cells within the theta cycle were not affected.

During MS cooling, LFP theta frequency decreased by 12 % and within-field firing rates of place cells decreased by a similar proportion (12%; interneurons 14%). This can explain why the fraction of place cells and the number of spikes within theta cycles were largely unaffected by MS manipulation. Cooling, therefore, exerted its main physiological effect by dispersing the same number of cell assemblies within the phase space of the prolonged theta cycles (Fig. 1b), as quantified by the longer τ in the face of preserved phase differences between place field peaks. Overall, these findings suggest that the fundamental organization of cell assemblies in the hippocampus is based on theta phase-preservation mechanisms, even at the expense of longer temporal lags between assemblies.

What are the mechanisms that keep the phase-organization within the MS-hippocampus loop when theta frequency changes? A previous experimental-computational model has shown that temporal lags between assemblies (τ) are correlated with the duration of the theta cycle (Geisler et al., 2010), but the physiological mechanism of the delays (τ) has remained undisclosed. One possibility is that the delays are a consequence of inhibition between competing successive cell assemblies. It was hypothesized previously that the mechanism responsible for the delays between cells assemblies may reside in hippocampal circuits (Geisler et al., 2010). However, the current findings demonstrate that the MS plays a critical role in adjusting the theta phase delays between neighboring place fields of pyramidal cells. Furthermore, following pharmacological inactivation of the MS, internally generated sequences disappear and theta-scale timing of place cells is impaired without affecting place cell positions in the maze (Wang et al., 2015). A putative mechanism may involve the reciprocal circuit between GABAergic neurons in MS and hippocampus. MS parvalbumin-expressing neurons innervate a variety of inhibitory neuron types in the hippocampus (Freund and Antal, 1988) and entorhinal cortex (Fuchs et al., 2016; Gonzalez-Sulser et al., 2014; Jeffery et al., 1995; Justus et al., 2017; Viney et al., 2018), while cholinergic neurons affect both principal cells and interneurons (Unal et al., 2015). In turn, long-range hippocampo-septal interneurons inhibit MS neurons (Gulyás et al., 2003; Takács et al., 2008; Toth et al., 1993). These hippocampo-septal inhibitory neurons are likely be of special importance because they are the best candidate for integrating the activity of multiple place cell assemblies and feedback population theta-oscillatory pulsing to subgroups of MS neurons.

### MS cooling-induced memory impairment

MS cooling exerted a robust effect on spatial memory, comparable to damage or pharmacologic inactivation of MS (Bolding et al., 2020; Chang and Gold, 2004; Chrobak et al., 1989; Givens and Olton, 1990; Winson, 1978). Choice errors correlated best with the temporal dynamic of MS cooling. Whether memory impairment is due to the documented physiological changes in the hippocampus or other mechanisms remains an open question.

In addition to MS neurons, our manipulation may have affected septal afferents, fibers of passage and possibly its surrounding structures. Therefore, one possible interpretation of the behavioral effects is that moderate cooling of LS, the n. accumbens and/or anterior end of the hippocampus is responsible for the behavioral deficit. However, lesion of LS does not have an impact on spatial behavior (Fraser et al., 1991; Galey et al., 1985; Leutgeb and Mizumori, 1999; Winson, 1978). Electrolytic lesions (Thifault et al., 1998) or ibotenic acid-induced lesions (Annett et al., 1989) of the nucleus accumbens did not impair spatial behavior and, in fact, damaging chatecholaminergic terminals selectively in the n. accumbens slightly improved spatial alternation (Taghzouti et al., 1985). Fibers of passage of subcortical neuromodulator neurons through the medial or lateral septum typically affect spontaneous alternation via the MS (Lalonde, 2002). Furthermore, memory errors reached maximum tens of seconds before mild cooling was observed in the hippocampus, suggesting that this mild secondary effect cannot explain memory impairment. Importantly, previous studies have shown that decreasing brain temperature even down to 30° C was not sufficient to impair spatial navigation in rats (Andersen and Moser, 1995; Moser and Andersen, 1994). In humans, cognitive functions are not affected until the brain is cooled to about 33°C (Fay and Smith, 1941). Thus, while we cannot exclude impaired spike transmission of fibers of passage in MS as a contributing factor, moderate cooling of the structures surrounding MS may not fully account for the observed severe memory deficit.

A remaining potential explanation is that the seemingly subtle but multiple changes of physiological parameters in the hippocampus (and expected but unobserved parallel changes in the entorhinal cortex) were responsible for the memory deficit. This possibility is supported by the observation that the temporal dynamic of choice errors correlated best with the time course of theta frequency decrease and that the maximum error rates occurred tens of seconds before a slight temperature decrease was detected in the hippocampus. Similarly, intra MS infusion of the tetracaine, muscimol and scopolamine suppressed theta oscillations and impaired performance in a spatial alternation task, and the choice errors best correlated with a change in theta oscillation frequency (Givens and Olton, 1990). During MS cooling, the same segment of the environment was ‘represented’ by the same number of neuronal assemblies but the time offsets between successive assemblies were longer in each theta cycle. Therefore, a possible explanation for the memory impairment, despite preserved spatial features of place cells, is that downstream reader structures, presumably not part of the MS-controlled theta system, failed to interpret the temporally altered hippocampal messages (Robbe and Buzsáki, 2009) but see (Venditto et al., 2019). Overall, our findings suggest that the fundamental organization scheme in the septo-hippocampal system is phase-based and that even minuscule temporal changes in large interconnected circuits may have behavioral consequences.

## Supplementary Material

**Supplementary Fig. 1.**
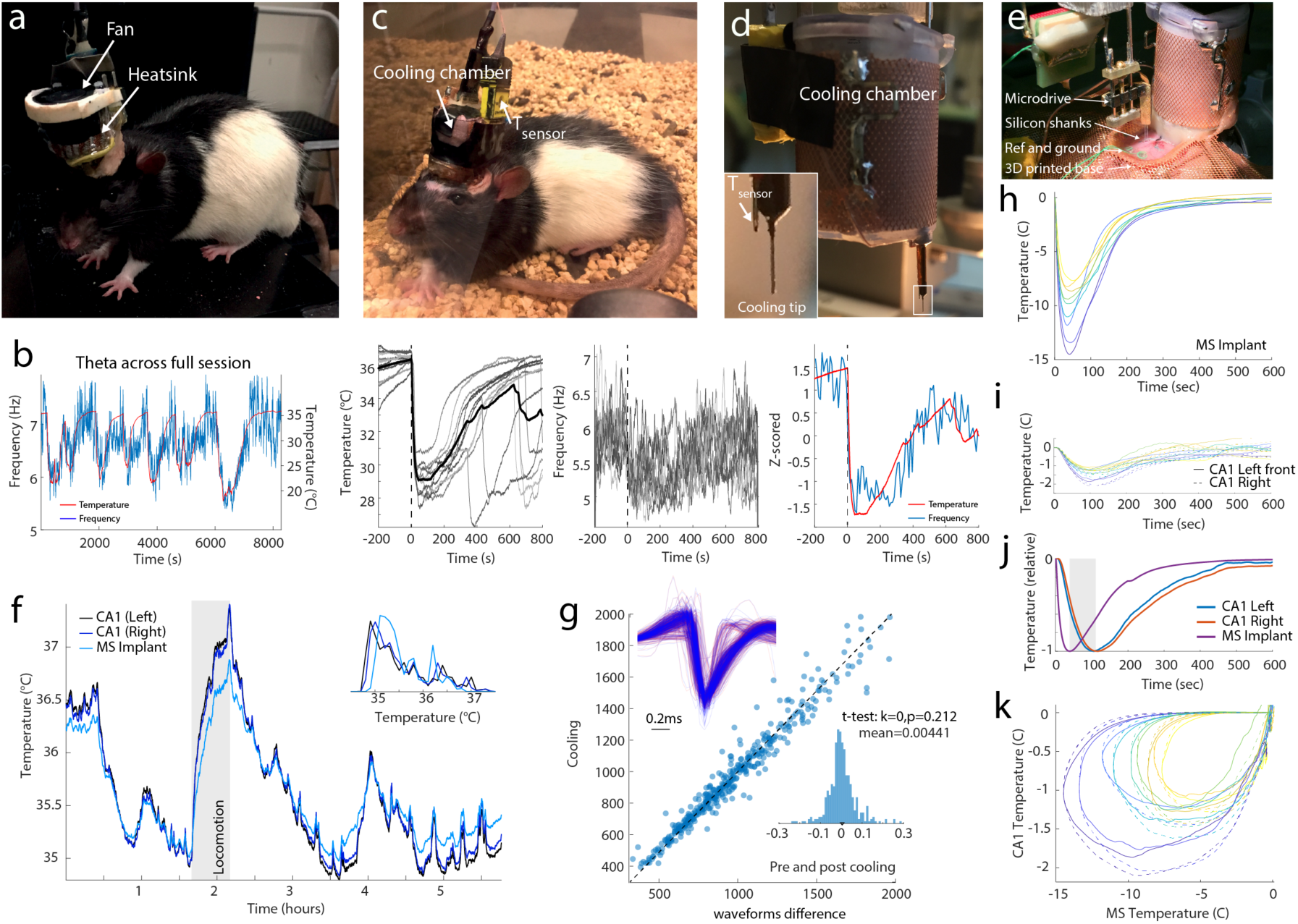
Related to Fig. 2. Methods of MS cooling. Two different methods for cooling the cryoprobe were tested. The first one used a Peltier device (panels a; Long and Fee, 2008). This allowed precise control of temperature over long duration and performing necessary tests of MS temperature control. However, it required both a heatsink and an electric fan for maintaining stable temperature of the Peltier element, increasing both volume and weight of the device. The second device was a small cooling reservoir (as also shown in Fig. 2a, b). This device was used in all physiological and behavioral experiments. **a**, Photograph of a Peltier device implanted in a rat. On top of the Peltier there is a small, low noise electric fan to cool the device. **b**, Relationship between MS local temperature (red) and theta frequency (blue) in the hippocampus. The MS was repeatedly cooling with dry ice. Second and third panels, successive cooling epochs are superimposed. Fourth panel, superimposed average MS temperature and frequency of hippocampal theta oscillations. Note reliable and non-accommodating relationship between MS temperature and hippocampal theta frequency across several hours. **c**, Photograph of a rat implanted with a dry iced-cooled cryoprobe. **d**, Details of the dry ice container and the cryoprobe with a temperature sensor connected to its outer tube. **e**, Photograph of the implantation with a silicon probe attached to a custom-built microdrive ready for implantation into CA1. Reference and ground screws and a 3D printed implant-base attached to the skull with meta-bond. **f**, Physiological temperature variation in the left and right hippocampus and MS. No cooling was induced in this session. Note that locomotion increases brain temperature (shaded column). Inset: Distribution of time windows spent in each temperature unit. Note >2° C span of temperature variation. **g**, Comparison of hippocampal CA1 unit waveforms before/after (red) and during (blue) MS cooling. Inset histogram shows that MS cooling did not significantly affect unit waveforms. **h**, Magnitude and time course of MS temperature and simultaneously monitored hippocampal temperature (**u**) in successive cooling sessions in a rat. **j**, Normalized temperature curves. Note the different temporal dynamics and > 50 s delay (shaded area) between the maximum cooling effects in MS and hippocampus. **k**, Lissajous orbit plots illustrate the temporal delays of temperature effects in the MS and hippocampus. Note that the temperature scale for the hippocampus (y) is ∼10 times smaller compared to the MS temperature scale (x).

**Supplementary Fig. 2.**
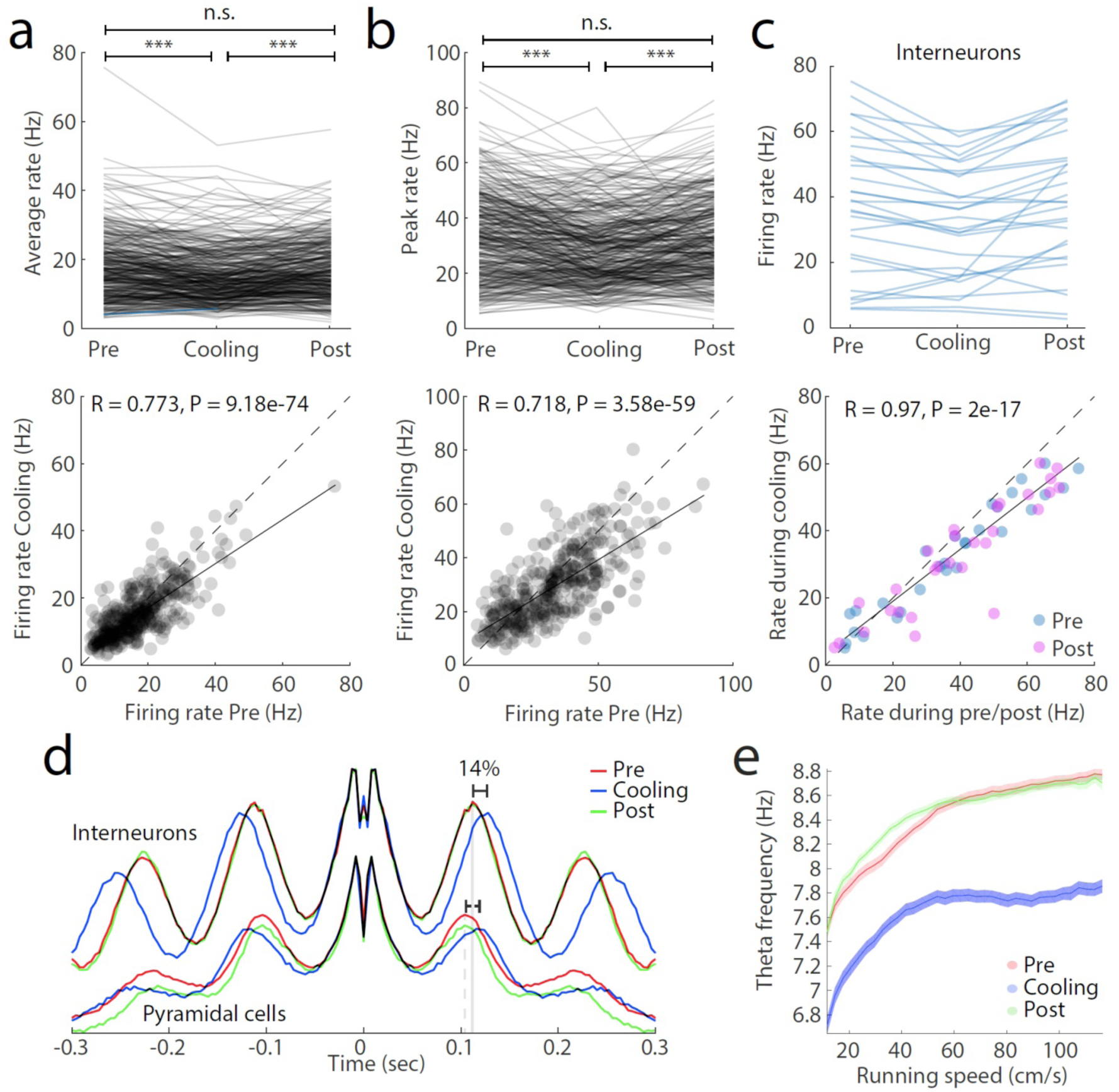
Related to Fig. 3. MS cooling affects single unit parameters. **a**, Firing rates of pyramidal neurons within their place fields (**a**) and their peak firing rates (**b**) (top) and illustration of the rate changes before and during MS cooling. R = 0.7, P < 10^−58^. **c**, Same display for putative fast firing interneurons. **d**, Autocorrelograms of putative interneurons and pyramidal cells before, during and after MS cooling. Note that pyramidal neurons oscillate faster than interneurons and the theta frequency oscillation of both interneurons and pyramidal cells decreases with MS cooling by approximately 14% on average. The autocorrelograms were constructed from all spikes occurring within place fields during running for pyramical cells, and across the whole maze for interneurons. **e**, Relationship between running speed and theta frequency as the animals runs through the circular maze, unlike in Figure 3g and h, where the values were obtained from individual place fields.

**Supplementary Fig. 3.**
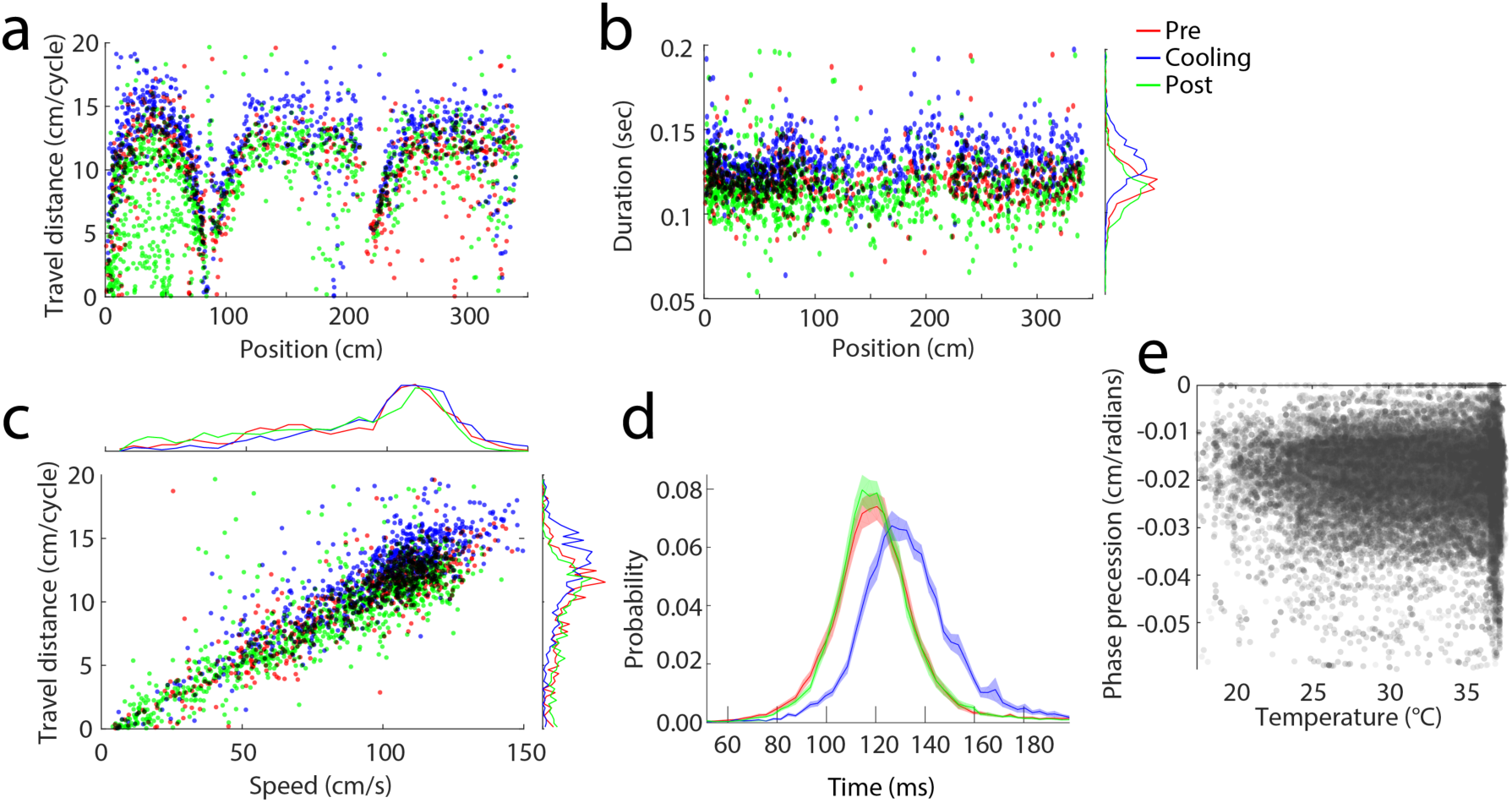
Related to Fig. 3. Speed and MS cooling affects are dissociable. **a**, Distribution of the travel distances per theta cycle during and after MS cooling as a function of the animal’s position on the track. 0-75 cm, center arm, followed by runs in the left or right arms of the maze. Note position-dependent variation of travel distances, due to speed variation on the track. **b**, Duration of the theta cycle as a function of the animal’s position. Note consistently longer theta duration during MS cooling. Side histograms show longer theta cycles during MS cooling. c. Distance traveled per theta cycle increases with running speed. Top histogram, speed distribution before, during and after MS cooling. This session was chosen for illustration purposes because running speed of the rat remained the same during the entire session. Side histograms show that longer distances are traveled per longer theta cycles during MS cooling. **d**, Distribution of theta cycle duration for all sessions (N = 21). Note that theta cycle duration increased during MS cooling but returned after cooling to the pre-cooling control duration, even though average speed gradually decreased from the beginning to the end of sessions (see Fig. 3e, Fig. 6c). **e**, Phase precession slope as a function of MS temperature. All values from before, during and after MS cooling trials are combined. Note low (R = -0.027) but significant (P = 7.67e-19) correlation.

**Supplementary Fig. 4.**
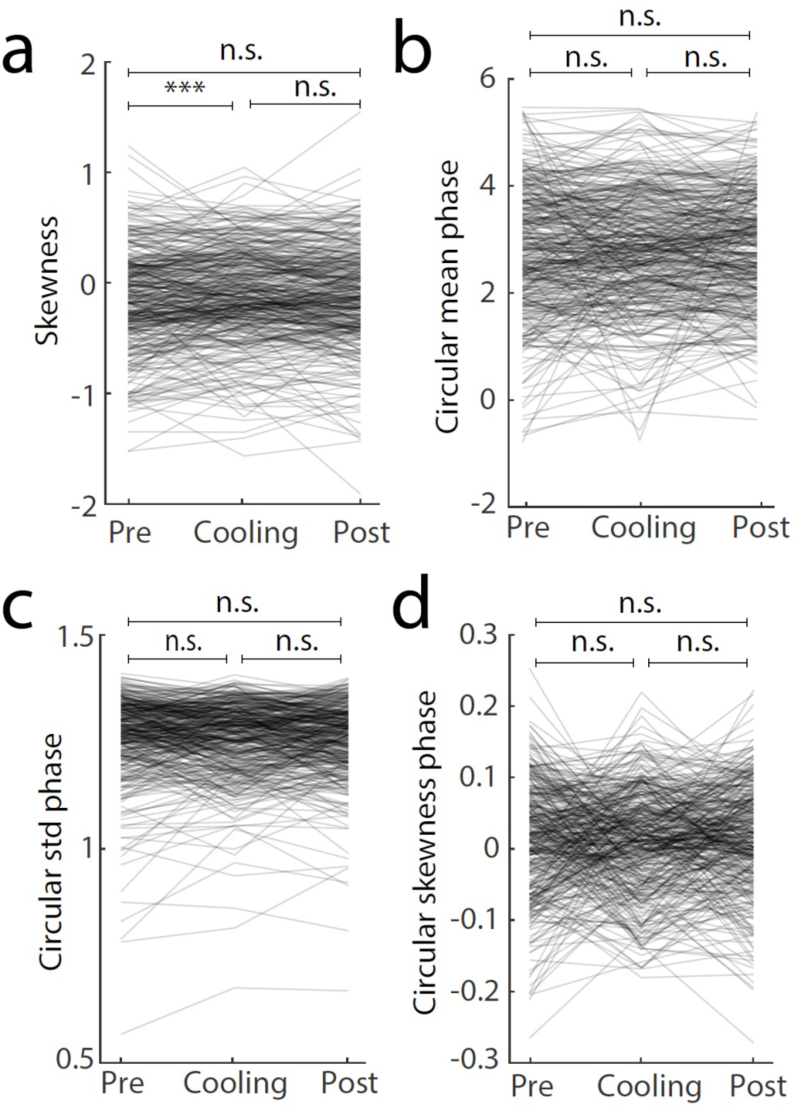
Related to Fig. 4. Properties of place fields during MS cooling. **a, b, c, d**, Skewness, mean phase (circular mean phase), standard deviation of phase (circular standard deviation), skewness phase (circular skewness), respectively, as a function of MS cooling.

**Supplementary Fig. 5.**
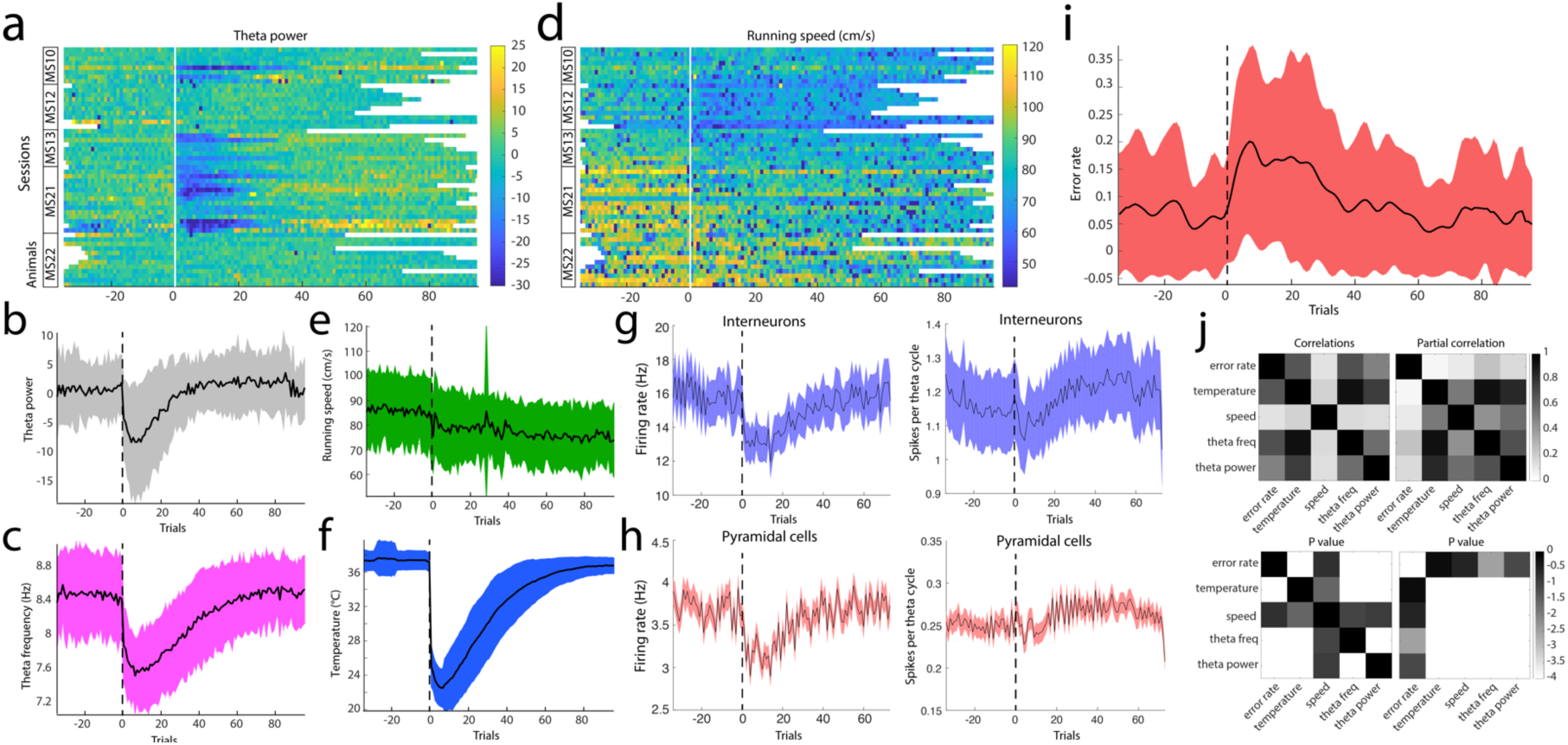
Related to Fig. 6. Time course of the effects of MS cooling on behavior and physiological parameters. **a, d**, Theta power and running speed as a function of MS cooling trials for each session. **b, e**, Group average and standard deviation related to panels **a** and **d**. Note monotonic deceleration of running speed over the entire session, likely reflecting the decreased motivation of the animal as it consumes more water cumulatively. **c, f, i**, Group means of theta frequency, MS temperature and choice errors, respectively. **g, h**, Trial-wise average firing rates for interneurons (**g**) and pyramidal cells (**h**) across 19 sessions with 319 interneurons and 1104 pyramidal cells. Right panels, spikes per theta cycle across the same populations. **j**, Correlation values (top) and P values (bottom) for each comparison (left panels). Partial correlation values (top) and P values (bottom) for each comparison (right panels). Note error rates correlate best with theta frequency and have no significant relationship to running speed.

## LEAD CONTACT AND MATERIALS AVAILABILITY

Further information and requests for resources and datasets should be directed to and will be fulfilled by György Buzsáki.

## Supplementary Material

### METHODS

#### Subjects and surgery

Rats (adult male Long-Evans, 250-450 g, 3-6 months old) were kept in a vivarium on a 12-hour light/dark cycle and were housed 2 per cage before surgery and individually after it. All experiments were approved by the Institutional Animal Care and Use Committee at New York University Medical Center.

Animals were anesthetized with isoflurane anesthesia and craniotomies were performed under stereotaxic guidance. A custom designed 3D printed cap (suppl. figure 1a) was attached to the skull with meta-bond, serving as a base for the probe implants and protection. A 12cm x 12cm sheet of copper mesh had beforehand been attached with dental cement to the base, from which a Faraday box/protector cap was later formed. Rats (Table 1) were implanted with silicon probes and tungsten wires to record local field potential (LFP) and spikes from the CA1 pyramidal layer (Vandecasteele et al., 2014). The tip of the cooling device was implanted at AP: +0.8mm, ML: 0.6mm (tilted 6° towards the midline), and lowered 6 mm below the brain surface, after which it was attached to the skull and base.

**Table 1:**
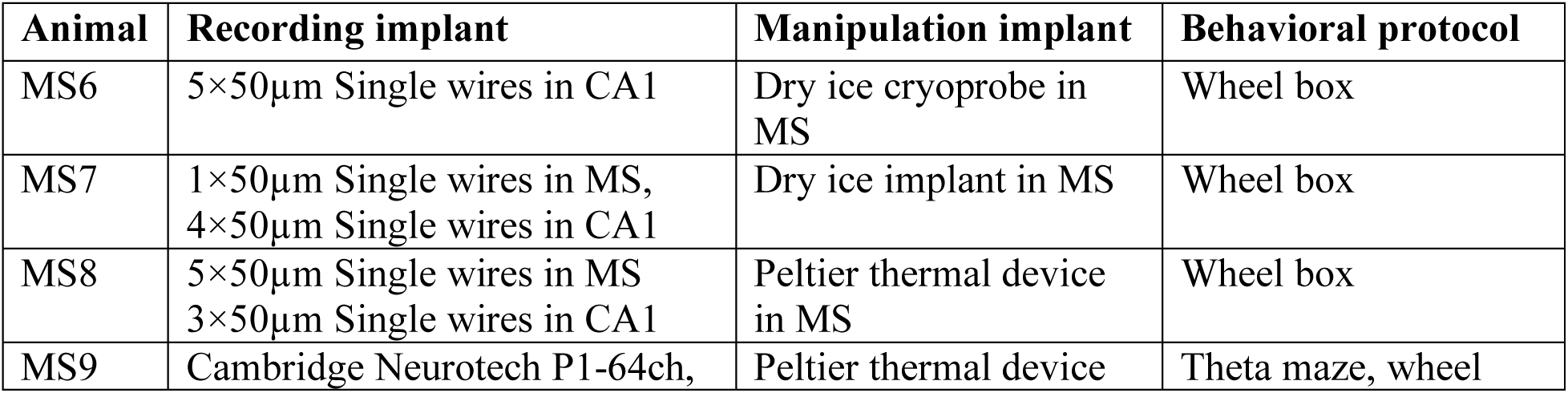
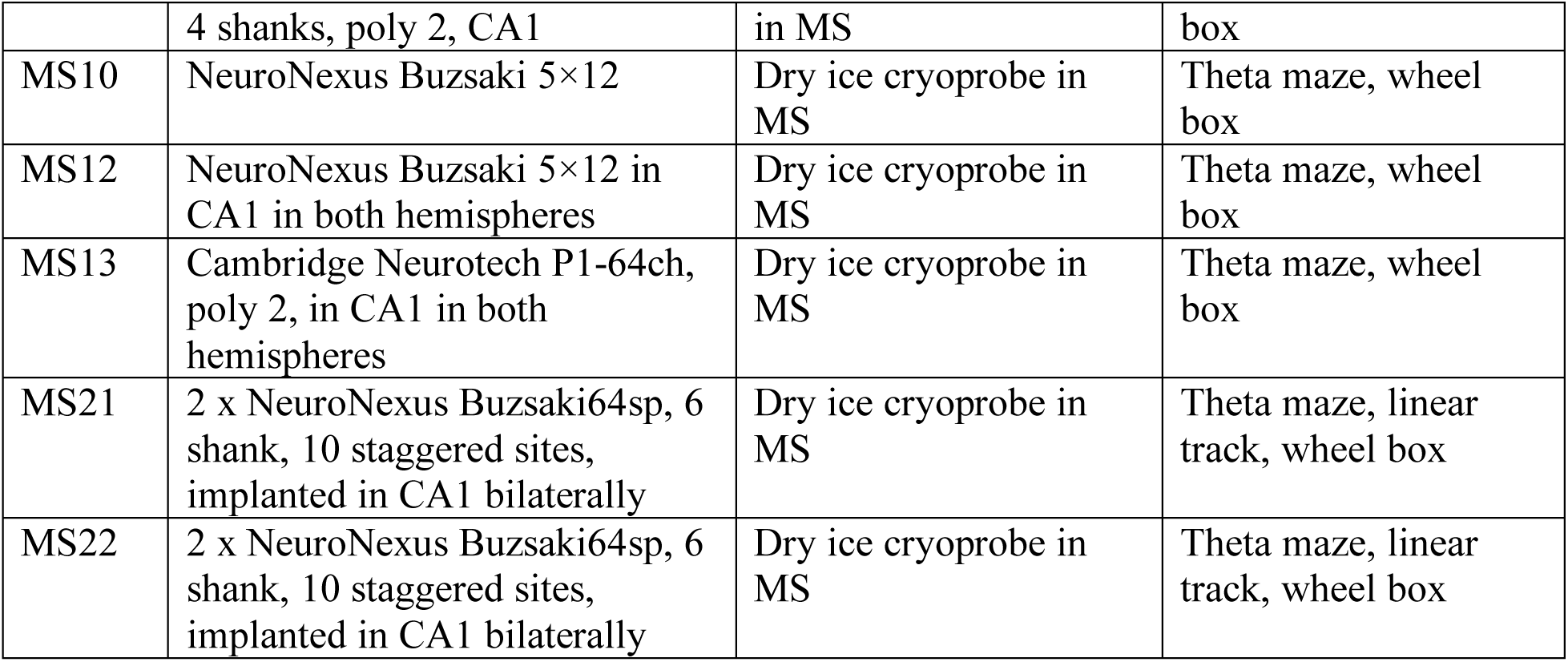
Animal implants and behavioral protocols

Silicon probes (NeuroNexus, Ann-Arbor, MI and Cambridge Neurotech, Cambridge, UK) were implanted in the dorsal hippocampus (antero-posterior (AP) -3.5mm from Bregma and 2.5 mm from the midline along the medial-lateral axis (ML)). The silicon probes consisted of three designs (table q): 4-shank with 16 sites per shank in a poly 3 staggered configuration, 5-shank with 12 sites each shank in a staggered configuration, and 6-shank probe with 10 sites per shank (Figure 2B). They were mounted on custom-made micro-drives (Suppl fig. 1e) to allow their precise vertical movement after implantation (Vandecasteele et al., 2012). Probes were implanted above the target region by attaching the micro-drives to the skull with dental cement (Suppl fig. 1e).

Craniotomies were sealed with sterile wax or gel. Stainless steel screws or 100µm steel wires were bilaterally screwed or implanted above the cerebellum, serving as ground and reference electrodes, respectively, for electrophysiological recordings (Suppl fig. 1e). At the end of electrode implantation and cryoprobe implantation (see below), the copper mesh was folded upwards, connected to the ground screw, and painted with dental cement. The mesh acts as a Faraday cage, shielding the recordings from environmental electric noise and muscle artifacts, provides structural stability and keep debris away from the probe implants. After post-surgery recovery, probes were moved gradually in 50 µm to 150 µm steps until they reached the CA1 the pyramidal layer. The pyramidal layer of the CA1 region was identified by physiological markers: increased unit activity, strong theta oscillations and phase reversal of the sharp wave ripple oscillations (Mizuseki et al., 2011).

#### Cooling devices

Two different cooling techniques were used. A Peltier device and dry ice in Styrofoam chamber (‘dry ice cryoprobe’). Both cooling devices were attached to a silver wire that conducted the cooling to the medial septum (Fig. 2a).

#### Dry ice cryoprobe

A 15mm diameter 3D printed container with lid was constructed and a hollow cylinder made from Styrofoam (outer diameter: 18mm, inner diameter: 10mm; thermal conductivity: 0.03 W/m) was inserted into the container. A 20 mm silver wire (127µm diameter, a-m systems, #781500; thermal conductivity: 406.0 W/mK), wrapped with graphene sheet (graphene-supermarket: Conductive Graphene Sheets, thickness: 25µm; thermal conductivity: 1300-1500W/m in x-y plane and 13-15W/m in z plane), was inserted through the base of the Styrofoam container and attached to the inside of the chamber with thermal adhesive (Arctic Silver Thermal Adhesive, ASTA-7G; thermal conductivity: >7.5W/m-K). 10mm of the wire was protruding from the base of the chamber. The protruding silver wire was then inserted into an 8 mm long hollow polyimide tube (1.1 mm diameter, Cole-Parmer 95820-09), such that 1.5mm of the silver wire was exposed. The polyimide tube was further sealed with adhesive in both ends to create a contained air insulation around the silver wire (figure 2A). The air insulation served as a thermal insulation (thermal conductivity: 0.024), to minimize the cooling effects along the wire (Aronov and Fee, 2011). Finally, a thermocouple (a temperature sensor: Omega, 80µm wires, product number 5SC-TT-K-40-72) was attached with epoxy adhesive to the cooling implant with the tip of the probe aligned with the protruding silver wire (figure 2A, and suppl. figure 1A). Cooling with dry ice was achieved by placing a small amount of dry ice into the “cooling chamber”, which conducted the cooling to the exposed implanted silver wire.

#### Peltier cooling device

The hot side of a two-stage Peltier device (custom thermoelectric, 04812-5L31-04CFG 2 Stage Thermoelectric/Peltier Module) was attached to a copper heatsink (5mm x 5 mm, Enzotech MOS-C10 Forged Copper MOSFET Heatsinks) with heat-conductive adhesive (Arctic Silver, Arctic Silver Thermal Adhesive). The heatsink was shaped to fit the inner dimensions of a 25 mm x 25 mm electric fan (GDSTIME, 5V DC Brushless fan). An 18 mm long silver wire (200µm diameter, a-m systems #782000) was attached to the cold side of the peltier device with heat conductive adhesive. An 8 mm long polyimide tube (1.1 mm diameter, Cole-Parmer 95820-09) was attached around the silver wire, sealed, and a thermocouple temperature sensor (Omega, 80µm wires, product number 5SC-TT-K-40-72) was attached to the tube. 1.5mm of the silver wire was exposed at the tip of the cooling device.

#### Electrophysiological Recordings

Animals were handled daily and accommodated to the experimenter before surgery. They were water restricted for 22 hours and trained to perform the behavioral task prior to surgery. After recovery from surgery, the animals were water restricted again to perform a spatial alternation task in a ‘theta’ (figure 8-shape) maze. The behavior session typically lasted 40 min, consisting of 40 control trials, after which the cooling was applied by manually placing a small amount of dry ice in the cooling chamber. The cooling typically peaked after about 60 sec after the cooling onset (suppl. fig. 1G) and lasted for about 10-12 minutes, corresponding to approximately 50 trials (figure 6C and suppl. Figure 2). The animal would continue the task for a total number of trials ranging from 80 to 200 (mean: 150 trials). The behavior was preceded and followed by 1-3 hours sleep sessions in the home cage of the rat.

#### Memory-demanding alternation task in a theta maze

In a ‘theta’ (figure 8-shape) maze (110 cm diameter, Fig. 2c), animals were trained to alternate between the left and the right arms to receive water drops at the reward locations (Figure 2C). The maze was placed on a platform 1 meter above the floor. The rat started from the reward location, ran along the central arm, after which it chose to run along the left or the right arm. If the animal performed the alternation correctly (visited the opposite arm than they visited in the previous trial), it received water reward. If it chose the wrong direction, the path to the reward location was blocked and the rat was forced to run back along the correct arm to collect reward.

The position of the animal was tracked with an OptiTrack 6-camera system (Natural Point Corp.). Calibration across cameras allowed for a three-dimensional reconstruction of the animal’s head position and orientation. A rigid body was created by mounting 6 reflective markers to a small 3D-printed holder, attached to the animal’s head-cap and tracked simultaneously by 6 infrared cameras (OptiTrack, Flex 3 cameras) at 120Hz.

## QUANTIFICATION AND STATISTICAL ANALYSIS

Electrophysiological recordings were conducted using an Intan recording system: RHD2000 interface board with Intan 64 channel preamplifiers sampled at 20 kHz (Intan Inc).

### Spike Sorting

Spike sorting was performed semi-automatically with KiloSort (Pachitariu et al., 2016) github.com/cortex-lab/KiloSort, using our own pipeline KilosortWrapper (a wrapper for KiloSort, github.com/petersenpeter/KilosortWrapper (Petersen et al., 2020), followed by a manual curation using the software Phy (github.com/kwikteam/phy) and our own designed plugins for phy (github.com/petersenpeter/phy-plugins).

### Unit Classification

Well isolated units were classified into putative cell types using the Cell Explorer petersenpeter.github.io/Cell-Explorer (Petersen and Buzsáki, 2020). Spiking characteristics, including the autocorrelograms, spike waveforms and putative monosynaptic connections derived from short-term cross-correlograms (English et al., 2017), were used to select and characterize well-isolated units. Three cell types were assigned: putative pyramidal cells, narrow and wide waveform interneurons.

### Theta phase and phase precession

An LFP channel located in CA1 was filtered in the theta range (4-10Hz, third order butter filter, filt-filt), and translated into phase by the Hilbert transform. The phase precession slope of a place field, was determined by performing a circular-linear regression of the position vs theta phase for all spikes within the boundaries of the place field.

### Oscillation Frequency of Neurons

For quantifying oscillation frequency of neurons, a 1ms-resolution spike raster was created and convoluted with an 80-point gaussian window. The auto-correlogram was calculated and the peak between 50ms and 150ms was determined and its reciprocal value was regarded as the oscillation frequency. For place field analysis (e.g. Fig. 3 B, C, F, H), only the spikes emitted as the animal passed through the field was included.

### Place cell analysis

Spiking data were binned into 3-5 cm wide segments of the maze, generating the raw maps of spike number and occupancy probability. Place field boundaries were manually defined by the following criteria: a spatial tuned firing rate, phase precession, and peak firing rate above 10Hz. For figure 5, place fields were automatically defined by the following criteria: at least 4 bins, where the firing rate was above 10% of the peak rate in the maze, peak firing rate > 8 Hz and spatial coherence > 0.6 (Hafting et al., 2008; Muller and Kubie, 1989).

### Spatial temporal compression

To determine the spatial-temporal compression, the distance between pairs of fields (determined as the spatial distance between the spatial peak firing rate) was plotted against the temporal delay inside theta cycles (either time or phase).

The temporal offset between individual overlapping fields was determined with a 1ms bin-sized cross correlogram, convoluted with a 60 bin-wide gaussian window. The phase offset was determined with a 0.01π bin-sized cross correlogram convoluted with a 60 bin-wide gaussian window. The compression and oscillation period were determined by fitting the surface equation below to the density of points (fig 5c):

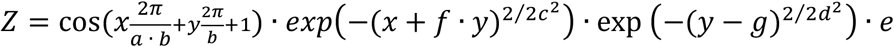

Where **a** is the compression (slope), **b** the CCG period (inverse of the oscillation frequency), **c** and **d** the widths of the two gaussian envelopes along the x and y dimensions, **e** the amplitude, **f** the x-y shift and **g** the y-offset.

### Statistical Analyses

All statistical analyses were performed with MATLAB functions or custom-made scripts. For rank order calculation, the probability of participation and firing rate correlations, the unit of analysis was single cells. Unless otherwise noted, for all tests, non-parametric two-tailed Wilcoxon rank-sum (equivalent to Mann-Whitney U-test), Wilcoxon signed-rank or Kruskal-Wallis one-way analysis of variance were used. Due to experimental design constraints, the experimenter was not blind to the manipulation performed during the experiment.

## DATA AND CODE AVAILABILITY

The dataset will be available from our data share via our website buzsakilab.com/wp/public-data/ (Petersen et al., 2018)

## KEY RESOURCES TABLE

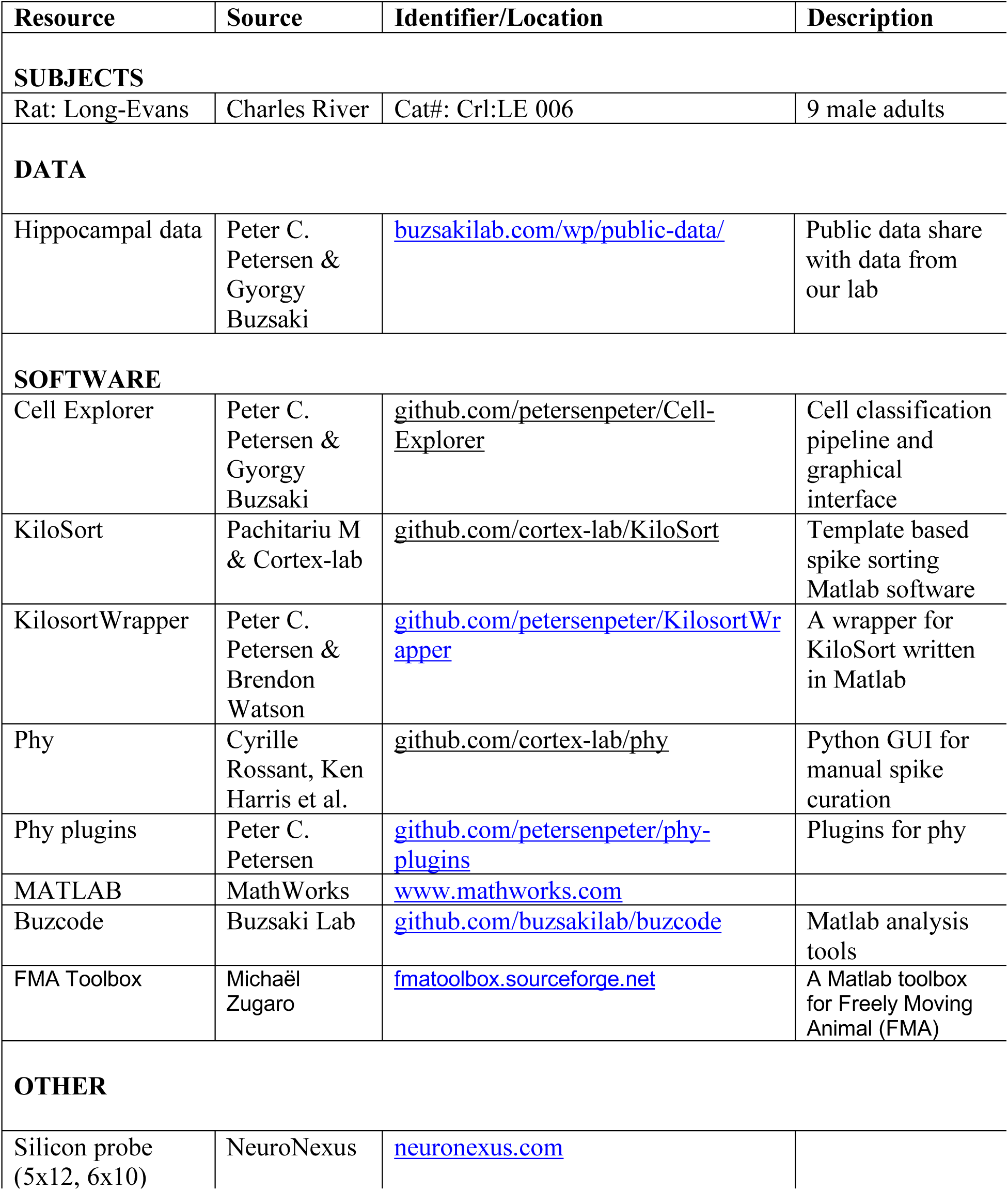
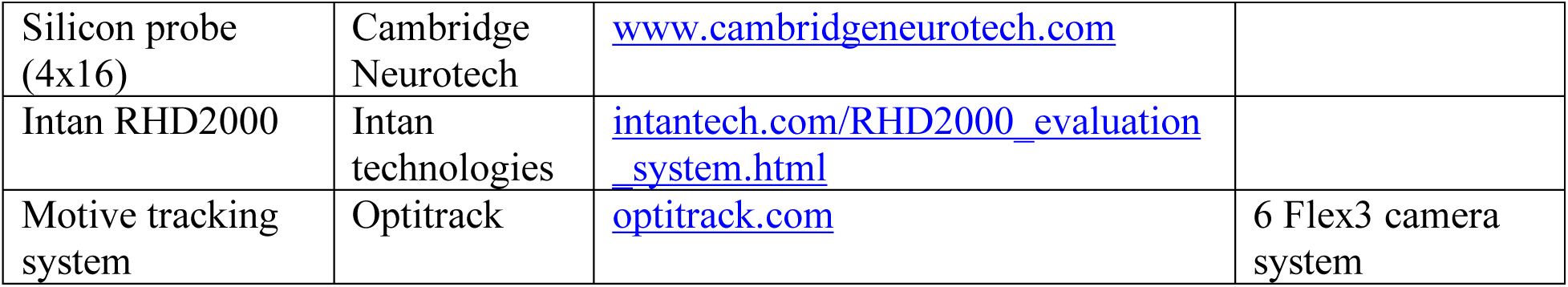

## Acknowledgements

We would like to thank Andrew Maurer, Viktor Varga, Manuel Valero and Antonio Fernandez-Ruiz for helpful comments on the manuscript. This work was supported by the Independent Research Fund Denmark, Lundbeckfonden Denmark, NIH, MH54671, MH107396, and NS 090583, NSF PIRE grant (#1545858), U19 NS107616 & U19 NS104590.

## Author Contributions

GB and PP designed the experiments. PP performed the experiments and analyzed the data. GB and PP wrote the paper.

## Reference list

Aghajan, Z.M., Acharya, L., Moore, J.J., Cushman, J.D., Vuong, C., and Mehta, M.R. (2015). Impaired spatial selectivity and intact phase precession in two-dimensional virtual reality. Nat. Neurosci. 18, 121–128.

Andersen, P., and Moser, E.I. (1995). Brain temperature and hippocampal function. Hippocampus 5, 491–498.

Annett, L.E., McGregor, A., and Robbins, T.W. (1989). The effects of ibotenic acid lesions of the nucleus accumbens on spatial learning and extinction in the rat. Behav. Brain Res. 31, 231–242.

Aronov, D., and Fee, M.S. (2011). Analyzing the dynamics of brain circuits with temperature: design and implementation of a miniature thermoelectric device. J. Neurosci. Methods 197, 32–47.

Bolding, K.A., Ferbinteanu, J., Fox, S.E., and Muller, R.U. (2020). Place cell firing cannot support navigation without intact septal circuits. Hippocampus 30, 175–191.

Borhegyi, Z. (2004). Phase Segregation of Medial Septal GABAergic Neurons during Hippocampal Theta Activity. J. Neurosci. 24, 8470–8479.

Brandon, M.P., Bogaard, A.R., Libby, C.P., Connerney, M.A., Gupta, K., and Hasselmo, M.E. (2011). Reduction of Theta Rhythm Dissociates Grid Cell Spatial Periodicity from Directional Tuning. Science 332, 595–599.

Burgess, N., Barry, C., and O’Keefe, J. (2007). An oscillatory interference model of grid cell firing. Hippocampus 17, 801–812.

Buzsáki, G. (2002). Theta Oscillations in the Hippocampus. Neuron 33, 325–340.

Buzsáki, G. (2010). Neural Syntax: Cell Assemblies, Synapsembles, and Readers. Neuron 68, 362–385.

Buzsáki, G., Czopf, J., Kondákor, I., and Kellényi, L. (1986). Laminar distribution of hippocampal rhythmic slow activity (RSA) in the behaving rat: Current-source density analysis, effects of urethane and atropine. Brain Res. 365, 125–137.

Chadwick, A., van Rossum, M.C., and Nolan, M.F. (2016). Flexible theta sequence compression mediated via phase precessing interneurons. ELife 5, e20349.

Chang, Q., and Gold, P.E. (2004). Impaired and spared cholinergic functions in the hippocampus after lesions of the medial septum/vertical limb of the diagonal band with 192 IgG-saporin. Hippocampus 14, 170–179.

Chrobak, J.J., Stackman, R.W., and Walsh, T.J. (1989). Intraseptal administration of muscimol produces dose-dependent memory impairments in the rat. Behav. Neural Biol. 52, 357–369.

Czurkó, A., Hirase, H., Csicsvari, J., and Buzsáki, G. (1999). Sustained activation of hippocampal pyramidal cells by ‘space clamping’ in a running wheel. Eur. J. Neurosci. 11, 344–352.

Dannenberg, H., Kelley, C., Hoyland, A., Monaghan, C.K., and Hasselmo, M.E. (2019). The firing rate speed code of entorhinal speed cells differs across behaviorally relevant time scales and does not depend on medial septum inputs. J. Neurosci.

Diba, K., and Buzsáki, G. (2008). Hippocampal Network Dynamics Constrain the Time Lag between Pyramidal Cells across Modified Environments. J. Neurosci. 28, 13448–13456.

Dragoi, G., and Buzsáki, G. (2006). Temporal Encoding of Place Sequences by Hippocampal Cell Assemblies. Neuron 50, 145–157.

Fay, T., and Smith, G.W. (1941). Observations on reflex responses during prolonged periods of human refrigeration. Arch. Neurol. Psychiatry 45, 215–222.

Fee, M.S., and Long, M.A. (2011). New methods for localizing and manipulating neuronal dynamics in behaving animals. Curr. Opin. Neurobiol. 21, 693–700.

Fraser, K.A., Poucet, B., Partlow, G., and Herrmann, T. (1991). Role of the medial and lateral septum in a variable goal spatial problem solving task. Physiol. Behav. 50, 739–744.

Freund, T.F., and Antal, M. (1988). GABA-containing neurons in the septum control inhibitory interneurons in the hippocampus. Nature 336, 170–173.

Fuchs, E.C., Neitz, A., Pinna, R., Melzer, S., Caputi, A., and Monyer, H. (2016). Local and Distant Input Controlling Excitation in Layer II of the Medial Entorhinal Cortex. Neuron 89, 194–208.

Fuhrmann, F., Justus, D., Sosulina, L., Kaneko, H., Beutel, T., Friedrichs, D., Schoch, S., Schwarz, M.K., Fuhrmann, M., and Remy, S. (2015). Locomotion, Theta Oscillations, and the Speed-Correlated Firing of Hippocampal Neurons Are Controlled by a Medial Septal Glutamatergic Circuit. Neuron 86, 1253–1264.

Galey, D., Durkin, T., Sifakis, G., Kempf, E., and Jaffard, R. (1985). Facilitation of spontaneous and learned spatial behaviours following 6-hydroxydopamine lesions of the lateral septum: a cholinergic hypothesis. Brain Res. 340, 171–174.

Geisler, C., Robbe, D., Zugaro, M., Sirota, A., and Buzsáki, G. (2007). Hippocampal place cell assemblies are speed-controlled oscillators. Proc. Natl. Acad. Sci. 104, 8149–8154.

Geisler, C., Diba, K., Pastalkova, E., Mizuseki, K., Royer, S., and Buzsáki, G. (2010). Temporal delays among place cells determine the frequency of population theta oscillations in the hippocampus. Proc. Natl. Acad. Sci. 107, 7957–7962.

Giocomo, L.M., Zilli, E.A., Fransén, E., and Hasselmo, M.E. (2007). Temporal Frequency of Subthreshold Oscillations Scales with Entorhinal Grid Cell Field Spacing. Science 315, 1719–1722.

Givens, B.S., and Olton, D.S. (1990). Cholinergic and GABAergic modulation of medial septal area: effect on working memory. Behav. Neurosci. 104, 849–855.

Gonzalez-Sulser, A., Parthier, D., Candela, A., McClure, C., Pastoll, H., Garden, D., Sürmeli, G., and Nolan, M.F. (2014). GABAergic Projections from the Medial Septum Selectively Inhibit Interneurons in the Medial Entorhinal Cortex. J. Neurosci. 34, 16739–16743.

Gothard, K.M., Skaggs, W.E., and McNaughton, B.L. (1996). Dynamics of Mismatch Correction in the Hippocampal Ensemble Code for Space: Interaction between Path Integration and Environmental Cues. J. Neurosci. 16, 8027–8040.

Gulyás, A.I., Hájos, N., Katona, I., and Freund, T.F. (2003). Interneurons are the local targets of hippocampal inhibitory cells which project to the medial septum. Eur. J. Neurosci. 17, 1861–1872.

Hafting, T., Fyhn, M., Bonnevie, T., Moser, M.-B., and Moser, E.I. (2008). Hippocampus-independent phase precession in entorhinal grid cells. Nature 453, 1248–1252.

Hangya, B., Borhegyi, Z., Szilágyi, N., Freund, T.F., and Varga, V. (2009). GABAergic Neurons of the Medial Septum Lead the Hippocampal Network during Theta Activity. J. Neurosci. 29, 8094–8102.

Harris, K.D., Henze, D.A., Hirase, H., Leinekugel, X., Dragoi, G., Czurkó, A., and Buzsáki, G. (2002). Spike train dynamics predicts theta-related phase precession in hippocampal pyramidal cells. Nature 417, 738–741.

Harris, K.D., Csicsvari, J., Hirase, H., Dragoi, G., and Buzsáki, G. (2003). Organization of cell assemblies in the hippocampus. Nature 424, 552–556.

Harvey, C.D., Collman, F., Dombeck, D.A., and Tank, D.W. (2009). Intracellular dynamics of hippocampal place cells during virtual navigation. Nature 461, 941–946.

Huxter, J., Burgess, N., and O’Keefe, J. (2003). Independent rate and temporal coding in hippocampal pyramidal cells. Nature 425, 828–832.

Jeffery, K.J., Donnett, J.G., and O’Keefe, J. (1995). Medial septal control of theta-correlated unit firing in the entorhinal cortex of awake rats. Neuroreport 6, 2166–2170.

Justus, D., Dalügge, D., Bothe, S., Fuhrmann, F., Hannes, C., Kaneko, H., Friedrichs, D., Sosulina, L., Schwarz, I., Elliott, D.A., et al. (2017). Glutamatergic synaptic integration of locomotion speed via septoentorhinal projections. Nat. Neurosci. 20, 16–19.

Kamondi, A., Acsády, L., Wang, X.J., and Buzsáki, G. (1998). Theta oscillations in somata and dendrites of hippocampal pyramidal cells in vivo: activity-dependent phase-precession of action potentials. Hippocampus 8, 244–261.

Katz, P.S., Sakurai, A., Clemens, S., and Davis, D. (2004). Cycle Period of a Network Oscillator Is Independent of Membrane Potential and Spiking Activity in Individual Central Pattern Generator Neurons. J. Neurophysiol. 92, 1904–1917.

Kay, K., Chung, J.E., Sosa, M., Schor, J.S., Karlsson, M.P., Larkin, M.C., Liu, D.F., and Frank, L.M. (2020). Constant Sub-second Cycling between Representations of Possible Futures in the Hippocampus. Cell 180, 552-567.e25.

Kjelstrup, K.B., Solstad, T., Brun, V.H., Hafting, T., Leutgeb, S., Witter, M.P., Moser, E.I., and Moser, M.-B. (2008). Finite Scale of Spatial Representation in the Hippocampus. Science 321, 140–143.

Lalonde, R. (2002). The neurobiological basis of spontaneous alternation. Neurosci. Biobehav. Rev. 26, 91–104.

Lee, M.G., Chrobak, J.J., Sik, A., Wiley, R.G., and Buzsáki, G. (1994). Hippocampal theta activity following selective lesion of the septal cholinergic systeM. Neuroscience 62, 1033–1047.

Leutgeb, S., and Mizumori, S.J.Y. (1999). Excitotoxic Septal Lesions Result in Spatial Memory Deficits and Altered Flexibility of Hippocampal Single-Unit Representations. J. Neurosci. 19, 6661–6672.

Lisman, J.E., and Jensen, O. (2013). The Theta-Gamma Neural Code. Neuron 77, 1002–1016.

Long, M.A., and Fee, M.S. (2008). Using temperature to analyse temporal dynamics in the songbird motor pathway. Nature 456, 189–194.

Lubenov, E.V., and Siapas, A.G. (2009). Hippocampal theta oscillations are travelling waves. Nature 459, 534–539.

Maurer, A.P., Cowen, S.L., Burke, S.N., Barnes, C.A., and McNaughton, B.L. (2006). Organization of hippocampal cell assemblies based on theta phase precession. Hippocampus 16, 785–794.

Maurer, A.P., Burke, S.N., Lipa, P., Skaggs, W.E., and Barnes, C.A. (2012). Greater running speeds result in altered hippocampal phase sequence dynamics. Hippocampus 22, 737–747.

McNaughton, B.L., Barnes, C.A., and O’Keefe, J. (1983). The contributions of position, direction, and velocity to single unit activity in the hippocampus of freely-moving rats. Exp. Brain Res. 52, 41–49.

McNaughton, N., Ruan, M., and Woodnorth, M.-A. (2006). Restoring theta-like rhythmicity in rats restores initial learning in the Morris water maze. Hippocampus 16, 1102–1110.

Mehta, M.R., Lee, A.K., and Wilson, M.A. (2002). Role of experience and oscillations in transforming a rate code into a temporal code. Nature 417, 741–746.

Mizuseki, K., Sirota, A., Pastalkova, E., and Buzsáki, G. (2009). Theta Oscillations Provide Temporal Windows for Local Circuit Computation in the Entorhinal-Hippocampal Loop. Neuron 64, 267–280.

Mizuseki, K., Diba, K., Pastalkova, E., and Buzsáki, G. (2011). Hippocampal CA1 pyramidal cells form functionally distinct sublayers. Nat. Neurosci. 14, 1174–1181.

Montgomery, S.M., Betancur, M.I., and Buzsáki, G. (2009). Behavior-Dependent Coordination of Multiple Theta Dipoles in the Hippocampus. J. Neurosci. 29, 1381–1394.

Moser, E.I., and Andersen, P. (1994). Conserved spatial learning in cooled rats in spite of slowing of dentate field potentials. J. Neurosci. 14, 4458–4466.

Muller, R.U., and Kubie, J.L. (1989). The firing of hippocampal place cells predicts the future position of freely moving rats. J. Neurosci. 9, 4101–4110.

O’Keefe, J., and Burgess, N. (2005). Dual phase and rate coding in hippocampal place cells: Theoretical significance and relationship to entorhinal grid cells. Hippocampus 15, 853–866.

O’Keefe, J., and Nadel, L. (1978). The Hippocampus as a Cognitive Map (Oxford: Clarendon Press).

O’Keefe, J., and Recce, M.L. (1993). Phase relationship between hippocampal place units and the EEG theta rhythm. Hippocampus 3, 317–330.

Pachitariu, M., Steinmetz, N.A., Kadir, S.N., Carandini, M., and Harris, K.D. (2016). Fast and accurate spike sorting of high-channel count probes with KiloSort. In Advances in Neural Information Processing Systems 29, D.D. Lee, M. Sugiyama, U.V. Luxburg, I. Guyon, and R. Garnett, eds. (Curran Associates, Inc.), pp. 4448–4456.

Pastalkova, E., Itskov, V., Amarasingham, A., and Buzsáki, G. (2008). Internally Generated Cell Assembly Sequences in the Rat Hippocampus. Science 321, 1322–1327.

Patel, J., Fujisawa, S., Berényi, A., Royer, S., and Buzsáki, G. (2012). Traveling Theta Waves along the Entire Septotemporal Axis of the Hippocampus. Neuron 75, 410–417.

Petersen, P.C., and Buzsáki, G. (2020). The Cell Explorer: a graphical user interface and a standardized pipeline for exploring and classifying single cells. https://petersenpeter.github.io/Cell-Explorer/ http://doi.org/10.5281/zenodo.3604173 (Zenodo).

Petersen, P.C., Hernandez, M., and Buzsáki, G. (2018). Public electrophysiological datasets collected in the Buzsaki Lab. https://buzsakilab.com/wp/public-data/ http://doi.org/10.5281/zenodo.3629881 (Zenodo).

Petersen, P.C., Watson, B., and Peyrache, A. (2020). KiloSortWrapper. https://github.com/petersenpeter/KilosortWrapper http://doi.org/10.5281/zenodo.3604165 (Zenodo).

Petsche, H., Stumpf, Ch., and Gogolak, G. (1962). The significance of the rabbit’s septum as a relay station between the midbrain and the hippocampus I. The control of hippocampus arousal activity by the septum cells. Electroencephalogr. Clin. Neurophysiol. 14, 202–211.

Ravassard, P., Kees, A., Willers, B., Ho, D., Aharoni, D., Cushman, J., Aghajan, Z.M., and Mehta, M.R. (2013). Multisensory Control of Hippocampal Spatiotemporal Selectivity. Science 340, 1342–1346.

Robbe, D., and Buzsáki, G. (2009). Alteration of Theta Timescale Dynamics of Hippocampal Place Cells by a Cannabinoid Is Associated with Memory Impairment. J. Neurosci. 29, 12597–12605.

Royer, S., Sirota, A., Patel, J., and Buzsáki, G. (2010). Distinct Representations and Theta Dynamics in Dorsal and Ventral Hippocampus. J. Neurosci. 30, 1777–1787.

Samsonovich, A., and McNaughton, B.L. (1997). Path Integration and Cognitive Mapping in a Continuous Attractor Neural Network Model. J. Neurosci. 17, 5900–5920.

Simon, A.P., Poindessous-Jazat, F., Dutar, P., Epelbaum, J., and Bassant, M.-H. (2006). Firing Properties of Anatomically Identified Neurons in the Medial Septum of Anesthetized and Unanesthetized Restrained Rats. J. Neurosci. 26, 9038–9046.

Skaggs, W.E., McNaughton, B.L., Wilson, M.A., and Barnes, C.A. (1996). Theta phase precession in hippocampal neuronal populations and the compression of temporal sequences. Hippocampus 6, 149–172.

Stewart, M., and Fox, S.E. (1990). Do septal neurons pace the hippocampal theta rhythm? Trends Neurosci. 13, 163–169.

Sweeney, J.E., Lamour, Y., and Bassant, M.H. (1992). Arousal-dependent properties of medial septal neurons in the unanesthetized rat. Neuroscience 48, 353–362.

Taghzouti, K., Louilot, A., Herman, J.P., Le Moal, M., and Simon, H. (1985). Alternation behavior, spatial discrimination, and reversal disturbances following 6-hydroxydopamine lesions in the nucleus accumbens of the rat. Behav. Neural Biol. 44, 354–363.

Takács, V.T., Freund, T.F., and Gulyás, A.I. (2008). Types and synaptic connections of hippocampal inhibitory neurons reciprocally connected with the medial septum. Eur. J. Neurosci. 28, 148–164.

Thifault, S., Krémarik, P., and Lalonde, R. (1998). Effects of Bilateral Electrolytic Lesions of the Medial Nucleus Accumbens on Exploration and Spatial Learning. Arch. Physiol. Biochem. 106, 297–307.

Thompson, S.M., Masukawa, L.M., and Prince, D.A. (1985). Temperature dependence of intrinsic membrane properties and synaptic potentials in hippocampal CA1 neurons in vitro. J. Neurosci. 5, 817–824.

Toth, K., Borhegyi, Z., and Freund, T.F. (1993). Postsynaptic targets of GABAergic hippocampal neurons in the medial septum-diagonal band of broca complex. J. Neurosci. 13, 3712–3724.

Tsanov, M. (2017). Speed and Oscillations: Medial Septum Integration of Attention and Navigation. Front. Syst. Neurosci. 11.

Unal, G., Joshi, A., Viney, T.J., Kis, V., and Somogyi, P. (2015). Synaptic Targets of Medial Septal Projections in the Hippocampus and Extrahippocampal Cortices of the Mouse. J. Neurosci. Off. J. Soc. Neurosci. 35, 15812–15826.

Vandecasteele, M., M, S., Royer, S., Belluscio, M., Berényi, A., Diba, K., Fujisawa, S., Grosmark, A., Mao, D., Mizuseki, K., et al. (2012). Large-scale Recording of Neurons by Movable Silicon Probes in Behaving Rodents. JoVE J. Vis. Exp. e3568.

Vandecasteele, M., Varga, V., Berényi, A., Papp, E., Barthó, P., Venance, L., Freund, T.F., and Buzsáki, G. (2014). Optogenetic activation of septal cholinergic neurons suppresses sharp wave ripples and enhances theta oscillations in the hippocampus. Proc. Natl. Acad. Sci. 111, 13535–13540.

Venditto, S.J.C., Le, B., and Newman, E.L. (2019). Place cell assemblies remain intact, despite reduced phase precession, after cholinergic disruption. Hippocampus 29, 1075–1090.

Vertes, R.P., and Kocsis, B. (1997). Brainstem-diencephalo-septohippocampal systems controlling the theta rhythm of the hippocampus. Neuroscience 81, 893–926.

Viney, T.J., Salib, M., Joshi, A., Unal, G., Berry, N., and Somogyi, P. (2018). Shared rhythmic subcortical GABAergic input to the entorhinal cortex and presubiculum. ELife 7, e34395.

Volgushev, M., Vidyasagar, T.R., Chistiakova, M., Yousef, T., and Eysel, U.T. (2000). Membrane properties and spike generation in rat visual cortical cells during reversible cooling. J. Physiol. 522, 59–76.

Wang, Y., Romani, S., Lustig, B., Leonardo, A., and Pastalkova, E. (2015). Theta sequences are essential for internally generated hippocampal firing fields. Nat. Neurosci. 18, 282–288.

Winson, J. (1978). Loss of hippocampal theta rhythm results in spatial memory deficit in the rat. Science 201, 160–163.

Zutshi, I., Brandon, M.P., Fu, M.L., Donegan, M.L., Leutgeb, J.K., and Leutgeb, S. (2018). Hippocampal Neural Circuits Respond to Optogenetic Pacing of Theta Frequencies by Generating Accelerated Oscillation Frequencies. Curr. Biol. 28, 1179-.e3.

